# A Genetically Engineered Human Organoid Model Reveals Distinct Genetic and Epigenetic Barriers of Lineage Plasticity in Early PDAC Transformation

**DOI:** 10.64898/2026.03.09.710586

**Authors:** Xian Zhang, Wilfred Wong, Hyein S. Cho, Dapeng Yang, Gary Dixon, Renhe Luo, Dingyu Liu, Denis Torre, Shigeaki Umeda, Federico González, Gokce Askan, Aslihan Yavas, Jeyaram Ravichandran Damodaran, Robert W. Rickert, Rajya Kappagantula, Fong Cheng Pan, Victoria G. Aveson, Adrien Grimont, Julian Pulecio, Samuel J. Kaplan, Heng Pan, Steven D. Leach, Christine A. Iacobuzio-Donahue, Rohit Chandwani, Christina S. Leslie, Danwei Huangfu

**Affiliations:** Developmental Biology Program, Memorial Sloan Kettering Cancer Center; New York, NY, 10065, USA; Weill Cornell Graduate School of Medical Sciences, Weill Cornell Medical College; 1300 York Avenue, New York, NY 10065, USA; Computational and Systems Biology Program, Memorial Sloan Kettering Cancer Center; New York, NY, USA; Tri-Institutional Training Program in Computational Biology and Medicine; New York, NY, USA; Louis V. Gerstner Jr. Graduate School of Biomedical Sciences, Memorial Sloan Kettering Cancer Center; New York, NY, 10065, USA; Human Oncology and Pathogenesis Program, Memorial Sloan Kettering Cancer Center; New York, NY 10065, USA; David M. Rubenstein Center for Pancreatic Cancer Research, Memorial Sloan Kettering Cancer Center; New York, NY 10065, USA; Department of Pathology, Memorial Sloan Kettering Cancer Center; New York, NY 10065, USA; Hepatopancreatobiliary Service, Department of Surgery, Memorial Sloan Kettering Cancer Center; New York, NY 10065, USA; Department of Surgery, Weill Cornell Medicine; New York, NY 10065, USA; Sandra and Edward Meyer Cancer Center, Weill Cornell Medicine; New York, NY 10065, USA; Department of Physiology and Biophysics, Englander Institute for Precision Medicine, Institute for Computational Biomedicine, Weill Cornell Medicine; 1300 York Avenue, New York, NY 10065, USA; Dartmouth Cancer Center, Dartmouth College; Hanover, NH 03755, USA; Department of Cell and Developmental Biology, Weill Cornell Medicine; New York, NY 10065, USA; Functional Oncogenomics Lab, Fundació Institut d’Investigació Sanitària Illes Balears, 07120, Palma, Balearic Islands, Spain; State Key Laboratory of Female Fertility Promotion, Center for Reproductive Medicine, Department of Obstetrics and Gynecology, Peking University Third Hospital; Beijing 100191, China

**Keywords:** pancreatic ductal adenocarcinoma (PDAC), pancreatic progenitor organoid (PO), gene editing, tumor suppressor gene (TSG), oncogenic KRAS, activator protein 1 (AP-1), *TET1* suppression, DNA methylation, chromatin remodeling, lineage plasticity, lineage restoration therapy

## Abstract

The lack of accurate, human-based models recapitulating early-stage pancreatic ductal adenocarcinoma (PDAC) has hindered therapeutic development. Using pluripotent stem cell-derived pancreatic progenitor organoids, we established a human PDAC model that faithfully reproduces the genetic, epigenetic, and transcriptomic trajectory of tumor initiation and progression *in vitro*, validated against clinical datasets and histopathology. We demonstrate that *CDKN2A* loss, nearly universal in patients but dispensable in mouse models, is essential for neoplastic transformation when combined with *KRAS* and *TP53* mutations, while *SMAD4* loss promotes tumor progression. Multi-omics profiling reveals epigenetic repression of pancreatic lineage program during PDAC initiation, alongside oncogenic AP-1-driven chromatin remodeling. Notably, we identify *TET1* suppression as a mechanistic link between oncogenic ERK signaling and the hypermethylation and silencing of essential pancreatic transcription factors. This model captures the genetic and epigenetic determinants of human PDAC, reveals antagonism between oncogenic and lineage restriction programs, and supports TET-based lineage restoration as a promising early intervention strategy for high-risk individuals.

## Introduction

The diagnosis of pancreatic ductal adenocarcinoma (PDAC) often occurs at advanced stages with a dismal prognosis for patients^1,2^. However, the mechanisms triggering the malignant transformation of human PDAC remain poorly understood, and the diagnostic features of early PDAC and its high-risk precursor lesions^3^ have not been fully characterized. Mutational profiling of clinical samples of pancreatic intraepithelial neoplasia (PanIN), the most common precursor lesion of PDAC, highlights the significance of this early disease stage for the accumulation of genetic and epigenetic alterations. Mutations or expression alterations in the four most altered genes in PDAC – *KRAS*, *CDKN2A*, *TP53*, and *SMAD4* – are all evident in PanIN^4^. Notably, oncogenic *KRAS* mutation and loss of protein product of the tumor suppressor gene (TSG) *p16/CDKN2A* are detected in low-grade PanIN, while additional TSG mutations in *TP53* and *SMAD4* are associated with the progression to high-grade PanIN and invasive PDAC^5–7^. The pervasive alterations observed in *KRAS*^8^, *p16/CDKN2A*^9^, and *TP53*^8^ in human PDAC strongly suggest that oncogenic *KRAS* signaling and TSG alterations coordinate in neoplastic transformation preceding the development of malignant PDAC from precursor lesions. Additionally, the inactivation of *SMAD4* is associated with advanced PDAC and metastasis in patients^8,10,11^.

The essentiality of genetic events in PDAC transformation is supported by findings from genetically engineered mouse models (GEMMs). The expression of oncogenic *Kras* alone often induces PanIN in GEMMs^12^, while most existing mouse models of PDAC rely on combined mutations in *Kras* and a TSG, which effectively recreate the progression from PanIN to invasive PDAC^13,14^. The accurate reproduction of histological features of (pre)neoplastic lesions of PDAC within these models reinforces the central role for GEMMs in PDAC research. However, there are notable differences between GEMMs and clinical observations. In comparison with the universal inactivation of *CDKN2A* in human PDAC, neither autochthonous PDAC initiated by mutant *Kras* in the mouse embryonic pancreas nor adult acinar and ductal cell-specific mouse PDAC depends on *CDKN2A* alteration for tumor initiation^12,15–17^, highlighting the discordance between animal models and human disease, and emphasizing the need for faithful human cell-based models. To establish a human cell-based experimental platform to model PDAC development, we focused on identifying a cell type that mirrors cells-of-origin in the clinical context of PDAC. Mature pancreatic epithelia, normally resistant to oncogenic transformation, regain susceptibility during injury-induced pancreatitis in adult mouse PDAC models^14,15^. This process involves the reactivation of the embryonic progenitor program in the inflamed pancreatic epithelia^14,18^. Clinical studies have identified pancreatitis as a major risk factor for PDAC in patients^19,20^. Therefore, we hypothesized that human pluripotent stem cell (hPSC)-derived pancreatic progenitors (PP2), which exhibit the embryonic progenitor program, could closely mimic the inflamed/metaplastic epithelium for modeling the onset of PDAC associated with genetic mutations. By developing a robust self-renewing pancreatic progenitor organoid (PO) platform incorporated with efficient gene editing strategies, we bridge the gap between mouse model and human disease to define the genetic and epigenetic barriers in early PDAC tumorigenesis.

## Results

### Single mutation of *CDKN2A* or *TP53* in combination with oncogenic *KRAS* in POs is insufficient to drive PDAC initiation

We generated PDX1^+^/NKX6.1^+^ PP2 from hPSCs utilizing a directed differentiation platform (Figure S1A,B). To maintain the cells for gene editing and subsequent molecular characterization, we explored protocols for expanding these progenitors^21–24^ and developed a 3D pancreatic progenitor organoid (PO) culture enabling the proliferation of these cells for over 20 weeks (Figure S1C). Throughout this period, POs maintained expression of early pancreatic progenitor (PP1) markers *PDX1* and *SOX9* but lost the expression of PP2 marker *NKX6.1* (Figure S1D,E).

To examine whether mutations of *CDKN2A* and *TP53* in combination with *KRAS* are sufficient to drive PDAC as shown in the mouse models, we established hPSCs capable of expressing a *GFP* tagged oncogenic *KRAS^G12V^* and a *Cas9* (for inducible TSG inactivation) in doxycycline-dependent manner (Figure 1A). As a control, we created an hPSC line expressing *GFP* and *Cas9* upon doxycycline treatment (Figure S2A,B). By transducing the *KRAS^G12V^/Cas9*-inducible hPSCs with lentiviruses to express sgRNAs targeting *CDKN2A* or *TP53*, we generated hPSCs ready to induce *KRAS^G12V^*expression and TSG deletion, designated as *K*, *KC*, and *KP* (*K:KRAS^G12V^*, C:*CDKN2A* deletion*, P:TP53* deletion, Figure 1A). Genetically modified hPSCs were then differentiated into PP2 at an efficiency comparable to wild-type cells (>90% PDX1^+^/NKX6.1^+^) and transferred to the PO culture followed by doxycycline treatment to induce the expression of oncogenic *KRAS* and the corresponding TSG loss (Figure 1A). Targeted CRISPR sequencing supported a high knockout rate (frameshift and large deletion>21bp) at *CDKN2A* and *TP53* loci in *KC* and *KP* POs 7 weeks after the treatment, confirmed by Western blotting (Figure 1B, S2D).

**Figure 1.**
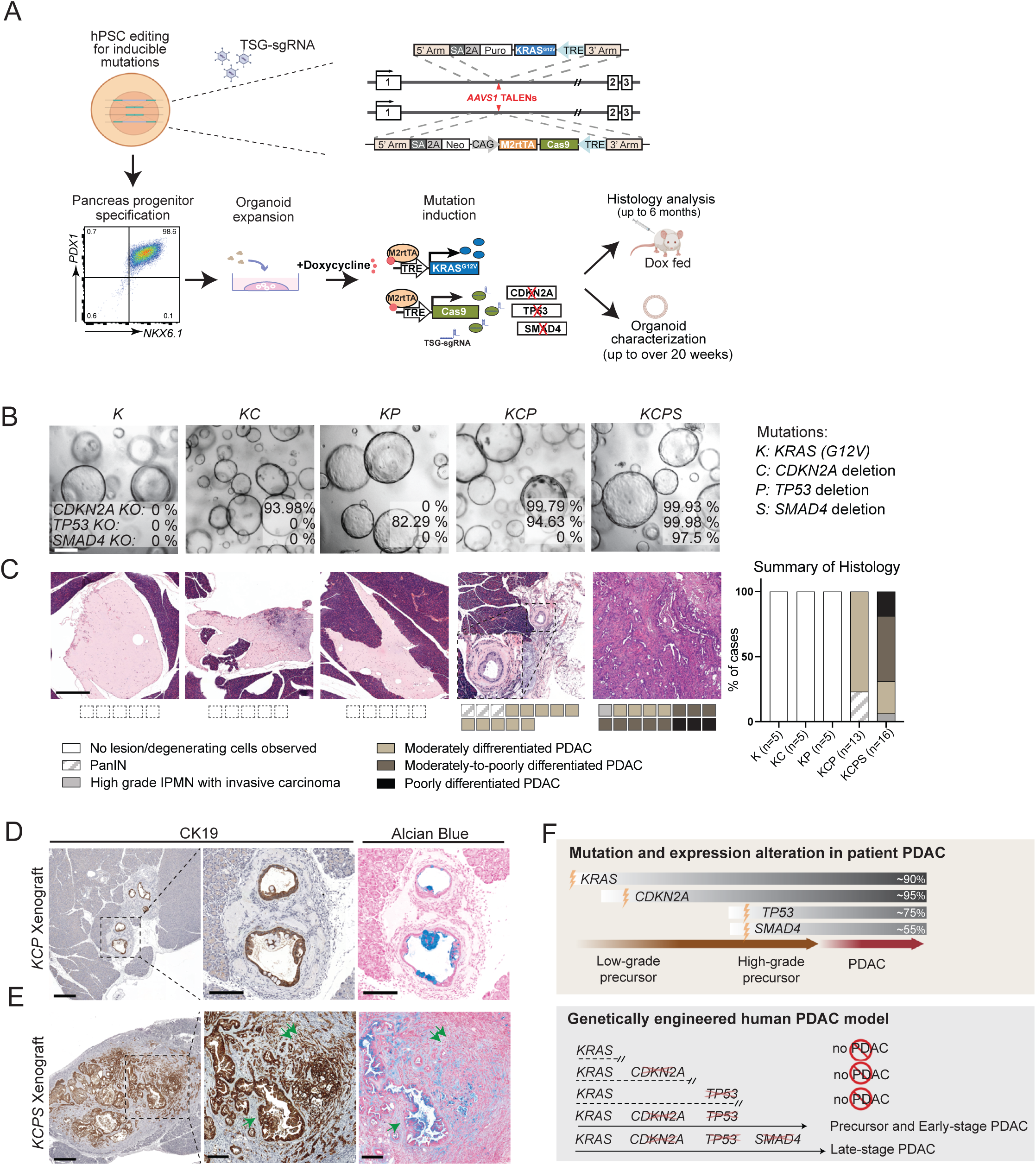
A pancreatic progenitor organoid based disease model recapitulates the genetic progression of human PDAC. (A) The summary of PDAC modeling strategy is presented. hPSCs were edited to enable doxycycline-inducible mutations associated with PDAC. *GFP*-*KRAS^G12V^*(or *GFP* as in a control line) and *Cas9* expression cassettes under TRE promoters were inserted into AAVS1 sites of both alleles. Successfully modified hPSC clones were transduced with lentivirus expressing sgRNAs targeting PDAC associated TSGs. Edited non-doxycycline treated hPSCs were directly differentiated into PP2 and cultured in Matrigel supplied with PO medium. Doxycycline was applied to induce oncogenic *KRAS* expression and TSG deletion. Genotype confirmed POs were orthotopically injected into NSG mice for tumorigenesis assay. (B) Targeted CRISPR sequencing was performed on *K*, *KC* and *KP* POs treated with doxycycline for 7 weeks. Scale bar: 200μm. (C) The tumorigenesis capacity of the POs was assessed by orthotopic transplantation. Mouse pancreata transplanted with *K*, *KC*, *KP*, *KCP* and *KCPS* POs were subjected to H&E staining. Serial sectioning was performed until either scar tissue was observed or the largest section plane was reached. The tumorigenesis capacity of the *KCP* and *KCPS* POs was confirmed by histopathological analysis. The case number of total transplanted animals and pathology evaluation based on H&E were indicated, scale bars: 500μm. (D,E) CK19 IHC and alcian blue staining were performed on *KCP* and *KCPS* PO xenograft. Single and double arrows indicated moderately and poorly differentiated components in *KCPS* xenograft. Representative of 5 tumors per genotype, scale bars: 500μm (left), 200μm (middle and right). (F) Summary schema of mutation profiles in precursor PanIN lesions and PDAC based on patient specimens(5-7) and xenograft outcomes from the genetically engineered human organoid model. The frequencies of gene alterations observed in PDAC as reported in Wilentz et al. (5) were indicated alongside each gene. Corresponding xenograft results from the engineered organoid model recapitulated the progressive mutational landscape observed in human disease.

To examine the tumorigenic capacity of gene-edited POs, we transplanted genotype-validated *K*, *KC* and *KP* POs orthotopically into NSG mice that were continuously fed doxycycline water to induce mutant *KRAS* (Figure 1A). Despite the expression of oncogenic *KRAS*, no epithelial lesions were detected in animals injected with *K*, *KC*, and *KP* POs. Instead, scar reaction and dystrophic calcification of degenerating cells (green arrows) were occasionally observed, indicating these cells could not appropriately engraft (n=5 per genotype, Figure S2E). Thus, in contrast to the mouse models^25,26^, oncogenic *KRAS* and loss of either *CDKN2A* or *TP53* were unable to recapitulate fully transformed PDAC in human context in our xenograft analysis.

### Dual mutations of *CDKN2A* and *TP53* in combination with oncogenic *KRAS* in POs recreate premalignant PanIN and PDAC upon orthotopic transplantation

The progression to PDAC in patients is marked by cumulative alterations in multiple TSGs^5,6^. Thus, we continued to create *KCP* POs using the lentiviral vector expressing *CDKN2A* and *TP53* sgRNAs in tandem. Additional *SMAD4* deletion was introduced by directly delivering *SMAD4* sgRNA into the *KCP* POs and selecting for TGF-β resistant POs (Figure S2C). Successful TSG knockout was confirmed through CRISPR sequencing and Western blotting (Figure 1B, S2D).

In contrast to the lack of tumorigenesis observed in mice injected with *K*, *KC* or *KP* POs, all 13 animals injected with *KCP* POs formed lesions in the mouse pancreas. Histopathological analysis confirmed that the microscopic glandular lesions from *KCP* POs resembled PanIN (n=3/13) and moderately differentiated PDAC (n=10/13) observed in patients (Figure 1C). In comparison, *KCPS* xenografts showed more robust tumor growth and desmoplastic reaction (Figure 1C). Among all the lesions assessed, most were moderately-to-poorly differentiated PDAC (n=8/16), and no PanIN was observed in any cases (n=0/16), indicating rapid disease progression (Figure 1C, Table S1).

To further characterize the lesions observed in *KCP* and *KCPS* xenografts, immunohistochemistry (IHC) was performed. CK19 staining revealed the presence of tumor cells in the xenografts (Figure 1D,E, left two panels), which were positive for E-cadherin, high in Ki67 and positive for GFP (from GFP-KRAS^G12V^ fusion), confirming the origin of the tumor epithelia (Figure S2F-H). Alcian blue staining was robust in the glandular lesions in *KCP* and *KCPS* xenografts, indicating the presence of acidic mucins, the signal of which weakened in poorly differentiated tumor cells losing glandular organization in *KCPS* xenografts (double vs single arrow, Figure 1D,E). This confirmed the progression of *KCPS* tumor towards higher grade^27^. Importantly, assessment of all the xenograft sections failed to detect teratomas, an issue that has previously hindered the characterization of tumor-specific histopathology in hPSC-derived disease models. Thus, the histology of the xenografts supported the fidelity of this progressive PDAC model, which aligned with the genetic progression of human PDAC^5–7^ (Figure 1F). It demonstrated that *CDKN2A* loss, dispensable in widely used mouse PDAC models, is essential for neoplastic transformation, while accumulation of mutations in *KRAS, CDKN2A* and *TP53,* frequently observed in high-grade PanIN and prevalent in PDAC^5,8,9,28^, are necessary to confer tumorigenic capacity in POs. The resulting lesions, without evident presence of poorly differentiated tumors 6 months post-transplantation, suggested that *KCP* POs represent an early stage in neoplastic transformation (Table S1). Moreover, the substantial growth and the presence of high-grade tumor cells in *KCPS* xenografts indicate disease progression associated with the additional *SMAD4* mutation^10,11^.

### *KCP* and *KCPS* POs exhibit distinct molecular signatures of malignant transformation and progression

The acquisition of tumorigenic capacity in *KCP* POs indicated the combination of these three genetic alterations conferred malignant features in POs. Indeed, several PanIN and PDAC markers, including *MUC5AC*, *TFF1*, *KLF4*, *S100P* and *ANXA10*^29^, were upregulated or ectopically expressed in *in vitro* cultured *KCP* POs, as shown by RT-qPCR and supported by positive staining of gastric MUC5AC in *KCP* POs (Figure 2A-B). To examine the global transcriptome changes as a consequence of driver PDAC mutations, we performed RNA-seq on the POs generated, and gene set enrichment analysis (GSEA) confirmed the activation of *KRAS* signaling in all samples compared to the GFP POs (Figure S3A,B)^30^. We also obtained the uniformly processed RNA-seq data for GTEx normal pancreas (V8, n=330) and TCGA pancreatic cancer (PAAD, n=183) samples^8,31,32^. To benchmark our PO dataset against the GTEx and TCGA data, we projected the PO data onto the principal components defined by the GTEx and TCGA samples. The *K*, *KC* and *KP* POs, shown to be non-tumorigenic in orthotopic transplantation assays, clustered away from tumorigenic *KCP* and *KCPS* POs in the PCA plot. This separation was delineated by a hyperplane defined by a logistic regression model trained with differentially expressed genes between TCGA PAAD tumors and GTEx normal samples (Figure 2C). For a direct comparison of paired tumor/normal pancreas specimens, we selected patient PDAC microarray data from published studies (GSE15471, GSE28735, GSE62452) as independent clinical references, and performed a meta-analysis. GSEA confirmed that top upregulated genes in *KCP* vs *GFP* POs were also overrepresented in genes upregulated in the patient datasets (NES=2.37, FDR=7x10^-12^), while the most prominent downregulated genes identified in *KCP* POs were also downregulated in patients (NES=-1.67, FDR=3x10^-4^, Figure 2D, Table S2). Therefore, the acquisition of tumorigenesis capacity in *KCP* and *KCPS* POs aligns with a clinically confirmed malignant molecular signature.

**Figure 2.**
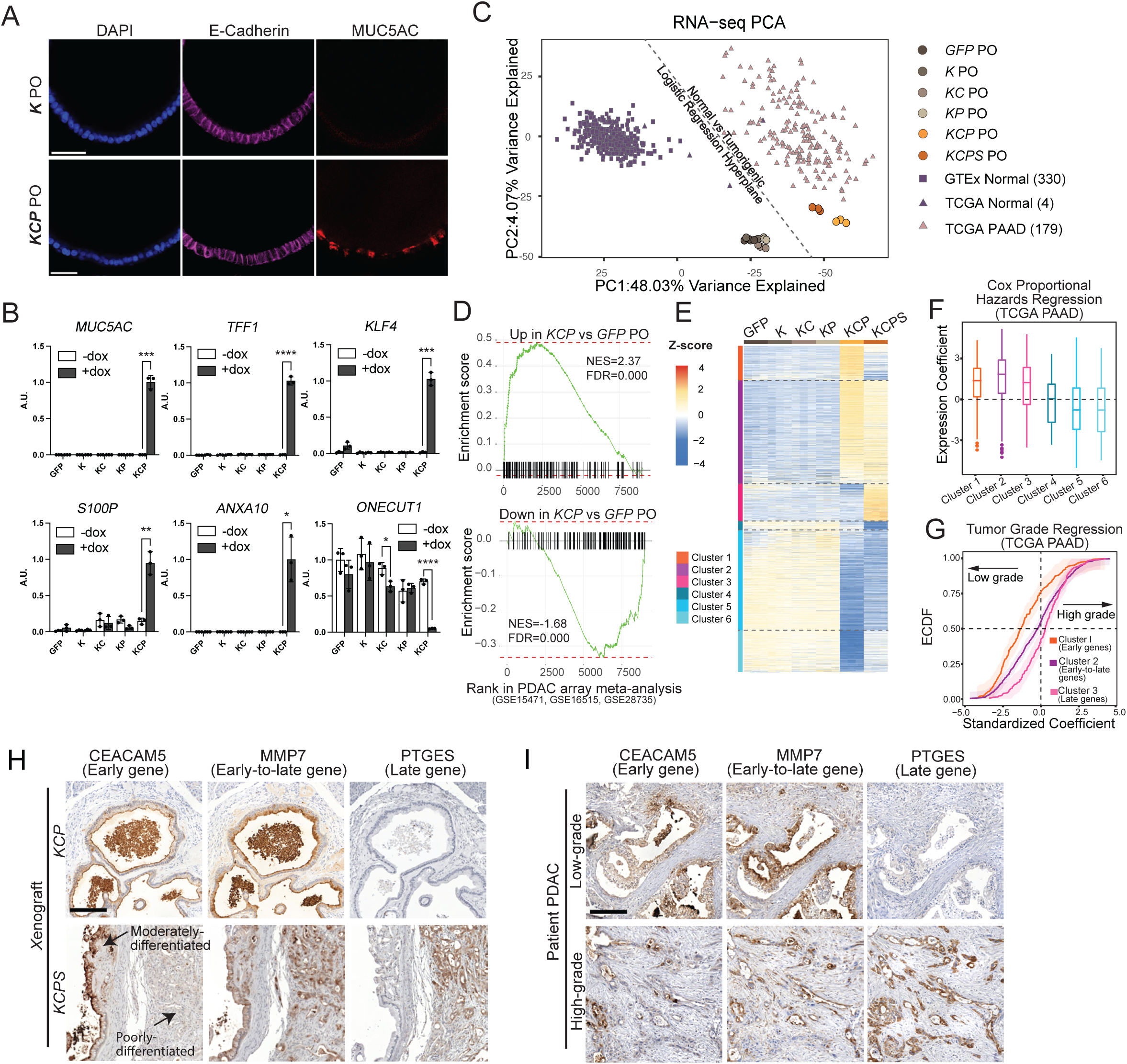
Transcriptome analysis identifies molecular signature associated with PDAC transformation and progression, validated by clinical data. (A) Immunofluorescence staining of E-cadherin and MUC5AC in POs was visualized by confocal microscopy and representative mid-plane section images were presented. Scale bars: 50μm. (B) RNA of passage matched POs (p20∼22) treated with or without doxycycline was collected and subjected to RT-qPCR analysis, mean±SD, n=3. Paired t-test, *Padj<0.05, **Padj<0.01, *** Padj<0.001, **** Padj<0.0001. (C) Principal component analysis was performed to embed Recount3 processed RNA-seq data from 183 TCGA PAAD and 330 GTEx pancreas samples. A logistic regression model was fit to differentiate non-tumorigenic GTEx from tumorigenic TCGA samples and the decision boundary was indicated by the dashed line. Recount3 processed RNA-seq data from POs were projected onto the PCA plot. (D) Paired patient PDAC/normal sample microarray data from GSE15471, GSE16515 and GSE29735 were used in a meta-analysis to obtain gene rankings. GSEA analysis was conducted using defined gene sets of top upregulated (n=166) and downregulated (n=146) genes in *KCP* POs comparing to *GFP* POs. (E) Clustering of top 3000 variable genes identified in PO RNA-seq data. (F) A Cox regression model was used with the normalized expression of each gene in the TCGA PAAD samples within the gene clusters defined in Figure 2E. A box and whisker plot of coefficients of each gene in the designated clusters was shown. (G) The association between gene expression and tumor grades in the TCGA PAAD cohort was assessed by logistic regression. The empirical cumulative distribution function with 95% confidence bands (obtained with bootstrap) of the z-scores for the regression coefficients are plotted. (H,I) IHC was performed to examine the expression of early, early-to-late gene and late gene in *KCP* and *KCPS* xenograft (H) or in low and high-grade human PDAC sections (I). Representative of 5 tumors per genotype or tumor grade. Scale bars: 200μm.

Clustering analysis of the top 3,000 variable genes in POs delineated distinct transcriptome features associated with PDAC development (Figure 2E, Table S3). Notably, Clusters 1,2 and 3, which represented genes upregulated in *KCP*, in *KCP* and *KCPS*, or in *KCPS* POs only, demonstrated overall positive association with poor prognosis in PDAC patients, as indicated by a positive Cox regression coefficient in TCGA PAAD patients. Conversely, genes in Clusters 5 and 6, which were downregulated in *KCP* or both *KCP* and *KCPS* POs, exhibited association with a favorable prognosis in patients. Genes from Cluster 4 (downregulated in *KCPS* PO only) did not show an association in either direction, likely due to the small size of the gene set (Figure 2F). Logistic regression to model the association of expression of genes in these clusters with tumor grades in TCGA PAAD led to clear stratification of Cluster 1 and Cluster 3, indicating their significant association with low and high-grade tumors, respectively (median coeff_grade_ 95% CI [-1.59, -1.08] and [0.09,0.44]). This aligned with their maximal expression in *KCP* or *KCPS* POs representing early and late stages of PDAC. The expression of Cluster 2 genes showed a weaker association with tumor grades (median coeff_grade_ 95% CI [-0.41, -0.04]), indicating their expression in both low and high-grade tumors, which is consistent with their expression in both *KCP* and *KCPS* POs (Figure 2G). Thus, we designated Cluster 1 and Cluster 3 as early progression (early) and late progression (late) genes, respectively, and Cluster 2 genes as early-to-late genes, based on their expression pattern during PDAC development in our model (Figure 2E,G).

To verify the expression of selected early, early-to-late, and late genes, we performed IHC on sections of *KCP* and *KCPS* xenografts. Consistently, early genes *CEACAM5* and *TFF1* were strongly expressed in the moderately differentiated components in both *KCP* and *KCPS* xenografts but were drastically lost in poorly differentiated tumor cells in *KCPS* xenografts. In contrast, staining of late genes, *PTGES* and *TRIM29*, preferentially labeled poorly differentiated components in *KCPS* tumors. Signals of early-to-late genes, *MMP7* and *CEACAM6*, were detected in both moderately and poorly differentiated components in the xenografts (Figure 2H, Figure S3C). We further examined PDAC patient samples and found decreased staining of *CEACAM5* and *TFF1,* and enhanced staining of *PTGES* and *TRIM29* in high-grade PDAC compared to low-grade tumors in patient sections. *MMP7* and *CEACAM6* staining covered all glandular structures in low-grade tumors and most poorly differentiated components in high-grade tumors (Figure 2I, S3D). These findings demonstrated that *KCP* and *KCPS* POs correspond to distinct stages of disease development, representing early transformed and progressed PDAC, respectively.

### Chromatin accessibility profiling reveals two integral components of cellular plasticity during PDAC development

To explore the molecular mechanisms driving malignant transformation and progression of *KCP* and *KCPS* POs, we performed ATAC-seq analysis on POs. To assess epigenetic changes in patient samples, we also developed a strategy to perform ATAC-seq on purified nuclei from PDAC epithelia by sorting aneuploid/polyploid nuclei from our institutional repository of snap-frozen and Optimal Cutting Temperature (OCT) medium-embedded patient tumors, thus circumventing issues associated with low cellularity in bulk PDAC specimens (Figure 3A, Table S4). Reflecting our RNA-seq findings, the chromatin accessibility profiles of *KCP* and *KCPS* POs clustered well with the majority of patient PDAC samples (n=14) but were clearly distinct from those of non-tumorigenic POs (Figure 3B).

**Figure 3.**
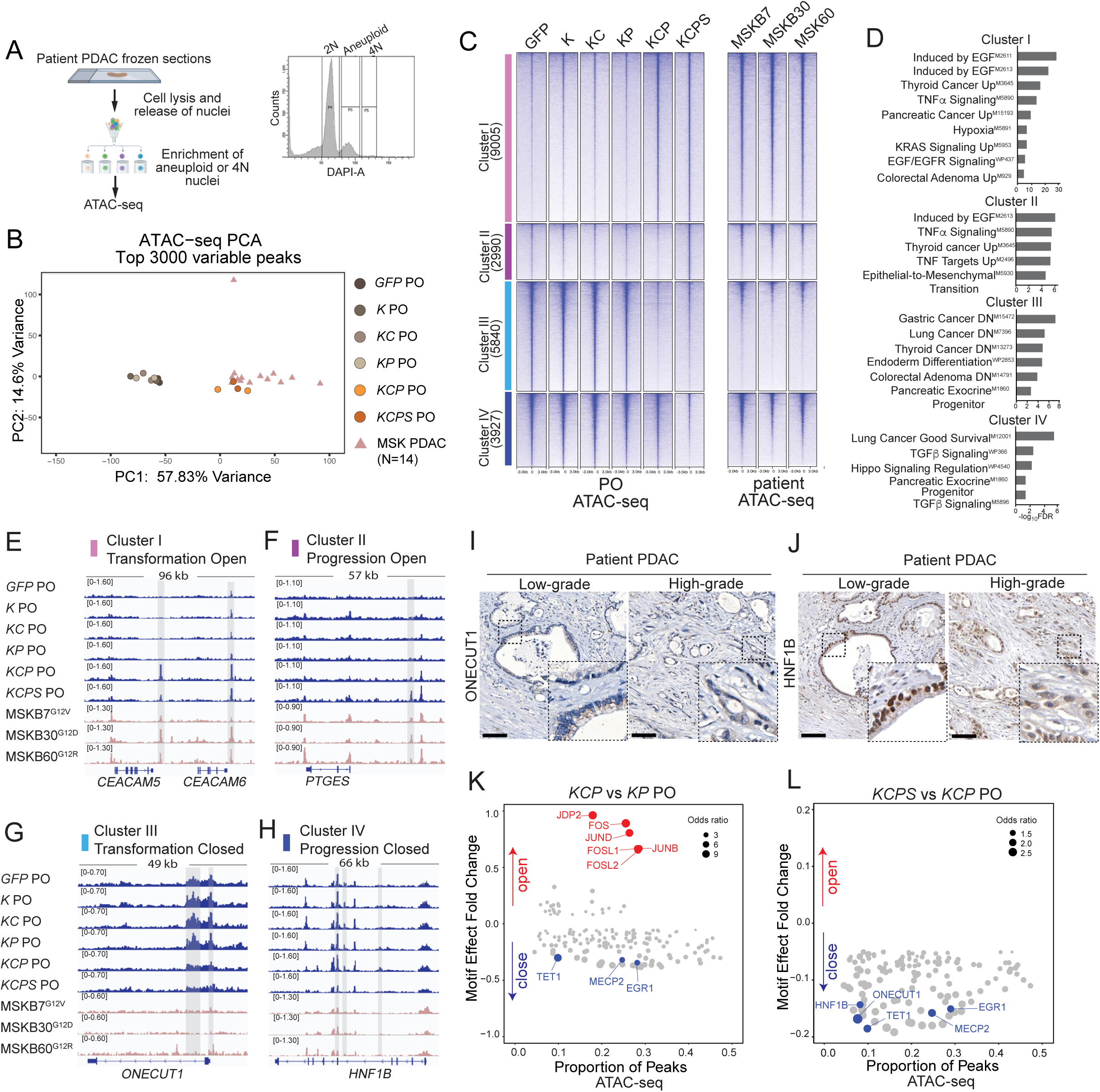
Chromatin accessibility profiling reveals two integral components of cellular plasticity during PDAC development. (A) Aneuploid or 4N polyploid nuclei from patient frozen sections were collected for chromatin accessibility assay. A representative flowcytometry plot was presented. (B) Omni-ATAC-seq was conducted on POs at matching passages in Figure 2C, n=2 per genotype. Patient PDAC ATAC-seq was performed on aneuploid or polyploid nuclei isolated (n=14). A PCA plot based on top variable regions in both POs and PDAC patients were generated. (C) Clustering of 21762 variable ATAC-seq peaks identified 4 clusters of regions with distinct behaviors in PDAC development in POs. ATAC-seq signals from corresponding genomic regions in selected patient samples were also presented. (D) MSigDB gene set discovery was conducted on each of the clusters in Figure 3C. Each ATAC-seq peak was assigned to the gene nearest to it. (E-H) Representative loci from the clusters in Figure 3C were visualized. Highlighted regions indicated identified peaks in distinct clusters in POs. (I,J) IHC analysis of *ONECUT1* and *HNF1B* expression in low and high-grade patient PDAC. Scale bars: 200μm. (K,L) Motif analysis was conducted to compare *KCP* PO with untransformed *KP* PO and *KCPS* with *KCP* PO. X-axis: the proportion of regions containing the motif; Y-axis: the median fold change of ATAC-seq signal associated with peaks containing or not containing the motif. The log odds ratio of the proportion of significantly open (or closed) peaks in the presence or absence of a specific motif was indicated in its absolute value. Motifs with significantly differential ATAC-seq signals (Padj<0.01) and with odds ratios > 1 were presented.

We clustered the top 21,762 variable ATAC-seq regions identified in POs into four groups based on their accessibility pattern during PDAC development, which was also confirmed in the patient samples (Figure 3C, Table S5). Cluster I regions (9,005 regions, designated as transformation open) exhibited increased accessibility in both *KCP* and *KCPS* POs compared to untransformed POs. These chromatin regions were linked by proximity to genes involved in KRAS, EGFR and TNF-α signaling and overlapped with gene sets upregulated in pancreatic cancer as well as several other endoderm malignancies in the Molecular Signatures Database (MSigDB, Figure 3D). Notably, Cluster I encompassed enhancers and promoters of PanIN/PDAC markers *CEACAM5*/*6, MMP7, ITGA2, MUC5AC* and *KLF4,* which exhibited elevated accessibility in *KCP* and *KCPS* POs and patient PDAC compared to non-tumorigenic POs (Figure 3E, S4A). Cluster II (2,990 regions, progression open) comprised regions that acquired accessibility during the progression of *KCPS* POs, including those associated with late genes *PTGES* and *EPSTI,* which were expressed specifically in *KCPS* PO and were identified as unfavorable prognosis markers for PDAC (Figure 3F, S4B,E,F).

We also examined regions with decreased accessibility in *KCP* and *KCPS* POs. Cluster III regions (5,840 regions, transformation closed) exhibited decreased accessibility in both *KCP* and *KCPS* POs (Figure 3C). These regions were associated with genes involved in endoderm differentiation, including pancreatic lineage transcription factors (TFs) *ONECUT1, NR5A2, PDX1*, and they showed similar loss of accessibility in patient PDACs (Figure 3D,G, S4C). Cluster IV regions (3,927 regions, progression closed) demonstrated a *KCPS*-specific decrease in accessibility (Figure 3C). Regions in these clusters were associated with pancreatic lineage TFs *HNF1B, MNX1, HHEX* and TGFβ signaling pathway members including *TGFBR2* (Figure 3D,H, S4D). The prominent association of pancreatic lineage genes with Clusters III and IV regions suggested that PDAC progression involves a loss of chromatin features associated with the pancreatic lineage program. Supporting this notion, we found that pancreatic specification involves the opening of Cluster III/IV and closing of Cluster I/II chromatin regions when we analyzed our ATAC-seq data from hPSC pancreas differentiation^33^ (Figure S4G).

Stepwise loss of accessibility in Cluster III and IV regions correlated with the distinct stages of PDAC development. We thus examined the expression of TFs associated with Cluster III/IV regions by proximity in the xenograft samples and patient PDAC. IHC staining detected a decrease of ONECUT1 signals in tumor epithelia of *KCP* xenografts and loss of signal in *KCPS* xenografts, in contrast to the normal ducts (Figure S4K,L). Meanwhile, human PDAC tumors exhibited an absence of nuclear staining for ONECUT1 in both low and high-grade PDACs (Figure 3I). This indicated the early loss of ONECUT1 during the acquisition of malignant status, consistent with our findings in the gene expression profile of POs and TCGA’s PDAC samples (Figure 2B, S4H,I), as well as previous studies^34,35^. HNF1B signal, on the other hand, remained comparable to normal ducts in *KCP* xenografts, but reduced nuclear staining was specifically detected in poorly differentiated components in *KCPS* xenografts (Figure S4M,N). Similarly, low-grade patient PDAC exhibited strong and distinct nuclear staining of HNF1B, which diminished in high-grade PDAC, in agreement with patient gene expression profiles (Figure 3J, S4J). Thus, we revealed a progressive suppression of genomic regions associated with the pancreatic lineage program and a downregulation of pancreatic TFs expression during malignant transformation and PDAC progression in our model. In particular, the reduction in HNF1B level coincided with the progression of cancer cells to a poorly differentiated state in *KCPS* xenografts. This observation is consistent with staining results in patient sections, as well as the clinical findings linking the loss of nuclear staining of HNF1B to a poor post-resection prognosis in a subset of PDAC patients^36^. Collectively, these results demonstrate the concordance of the model with established biological features of human disease.

### The acquisition of lineage plasticity during early PDAC transformation requires the activation of pro-transforming AP-1 factors

The alterations in chromatin accessibility during PDAC development indicates two mechanisms driving malignant transformation: gain of ectopic accessibility in regions associated with trans-lineage/malignant lineage and loss of accessibility in regions related to the pancreatic lineage program (Figure 3C). AP-1 motifs were highly enriched in the regions gaining accessibility during early malignant transformation of *KCP* POs (Figure 3K, S4O,P). To identify the crucial AP-1 factors involved in the cellular transformation, we prioritized candidates that displayed increased expression in *KCP* POs and performed Cas9-based perturbation. *FOSL1*, slightly upregulated in *KCP* POs, was included as a positive control, as it is a recognized vulnerability in *KRAS*-driven pancreatic cancer and demonstrated to be indispensable for mouse PDAC tumorigenesis^18,37,38^. High-efficiency gene perturbation was confirmed by targeted CRISPR sequencing and Western blotting 4 weeks after the introduction of sgRNAs in *KCP* POs (Figure 4A, S5A). The sensitivity of this approach was validated by the observed proliferation defects in *FOSL1*-depleted *KCP* POs, which showed the strongest reduction of pHH3 staining. Among other AP-1 factors examined, *FOS*-depleted *KCP* POs also displayed a decrease in cell proliferation (Figure 4B, S5A).

**Figure 4.**
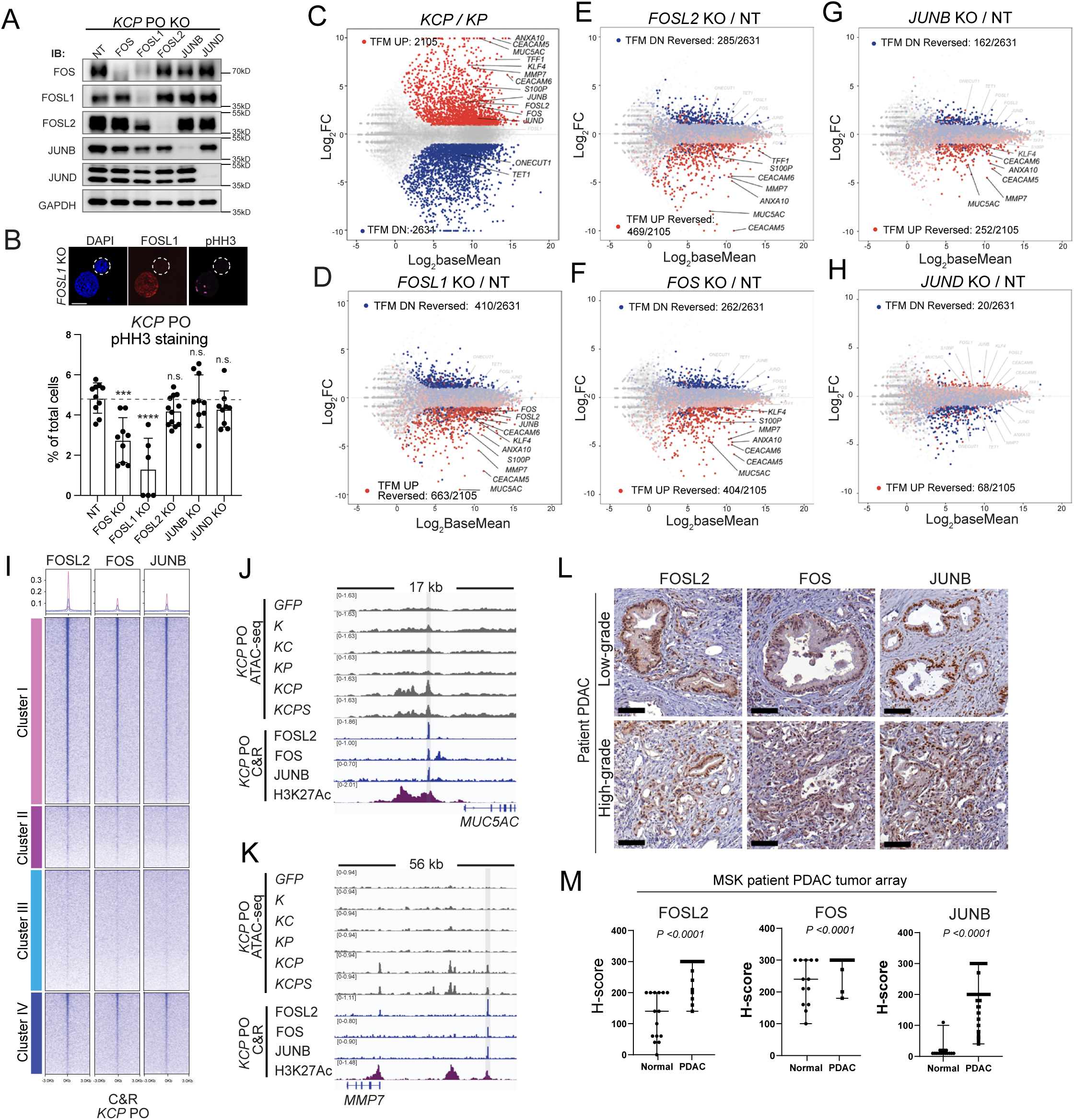
The acquisition of lineage plasticity during early PDAC transformation requires the activation of oncogenic AP-1 factors. (A) AP-1 perturbation screening was conducted in *KCP* POs at passage 8. Western blot analysis was performed four weeks after sgRNA introduction. (B) Immunofluorescence signal of pHH3 was visualized using confocal microscopy and percentage of pHH3 positive nuclei was quantified for each PO imaged. *** Padj<0.001, **** Padj<0.0001, one-way ANOVA. Immunofluorescence of *FOSL1* knockout POs were presented as an example. Scale bar: 100 μm. (C-H) TFM UP genes (n=2105, red, Log2FC>1, FDR<0.01) and TFM DN genes (n=2631, blue, Log2FC<-1, FDR<0.01) were defined by differentially expressed gene analysis comparing *KCP* and *KP* POs (c). For differential expression analysis in AP-1 depleted *KCP* POs, The TFM UP and DN genes were tracked on the MA plot, and the total number of reversed TFM UP and DN genes (|log2 FC| >1 and FDR <0.01), were annotated. (I) C&R assays were performed to identify FOSL2, FOS and JUNB bound genomic regions in *KCP* POs. A tornado plot illustrating the binding signal of these AP-1 targets was presented based on clusters identified in Figure 3C. Metapeaks represented the average of the signal in each cluster defined by the color. (J,K) Ectopically accessible regions associated with *MUC5AC* and *MMP7* genes in *KCP* POs were visualized alongside C&R tracks of FOSL2, FOS and JUNB. Statistically called genomics regions in C&R assays were highlighted. (L) IHC of FOSL2, FOS and JUNB was performed on patient sections of PDAC. Representative of 5 tumor samples each. Scale bars: 200μm. (M) Expression of *FOSL2*, *FOS* and *JUNB* was evaluated on the MSKCC’s PDAC tumor array (PDAC n=100, normal n=13). Mann Whitney U Test was performed on H-score of each AP-1 factor.

To evaluate the impact of individual AP-1 factors, we followed the behavior of transformation-associated upregulated and downregulated genes (TFM UP, n=2105; TFM DN, n=2631) identified in differentially expressed gene analysis of *KCP* vs *KP* POs (Figure 4C). As expected, *FOSL1* knockout decreased the amplitude of malignant transcriptome features as compared to the non-targeting sgRNA-transduced *KCP* PO (Figure 4D). *FOSL2*, *FOS* and *JUNB* depletion in *KCP* PO also led to a similar attenuation of malignant features, while *JUND* depletion had little impact on the transcriptome of *KCP* POs (Figure 4E-H, Table S6). RT-qPCR confirmed the significant downregulation of malignant markers *MUC5AC*, *CEACAM5*, *ANXA10*, *MMP7* upon *FOSL1*, *FOSL2*, *FOS* and *JUNB* knockout, while *S100P* and *KLF4* level exhibited a genotype specific downregulation in AP-1 knockout *KCP* POs. None of these genes showed downregulation in *JUND* knockout *KCP* POs (Figure S5B). The decrease of *MUC5AC* expression in *KCP* POs with *FOSL2*, *FOS* and *JUNB* deletion was verified via immunofluorescence, confirming a transforming role of these AP-1 factors in PDAC development (Figure S5C-E).

To determine how oncogenic AP-1 factors regulate the trans-lineage/malignant features, CUT&RUN (C&R) assays were conducted on *KCP* POs, and enriched binding of FOSL2, FOS, and JUNB was observed in Cluster I (transformation open) regions, with AP-1 motifs being predominantly detected (Figure 4I, S5F). Among those regions linked to trans-lineage/malignant markers, *MUC5AC* and *MMP7* were bound by the oncogenic AP-1 factors and labeled with the active histone marker H3K27ac in *KCP* POs, supporting a functional role for AP-1 in epigenetic reprogramming in PDAC (Figure 4J,K). IHC of the FOSL2, FOS and JUNB in the *KCP* and *KCPS* xenografts indicated enrichment of nuclear staining of these AP-1 factors in tumor epithelia (Figure S5H). Similarly, IHC analysis of a PDAC tumor array from MSKCC (PDAC: n=100 and normal pancreas: n=13) revealed a significant increase of these AP-1 factors in human PDAC samples (Figure 4L,M), in agreement with expression data of our POs and the TCGA PAAD (Figure S5G,I). Notably, consistent with its transforming role in *KCP* POs, *FOSL2* expression is associated with poor patient prognosis in the TCGA PAAD dataset. This contrasts with the favorable prognosis associated with the expression of *JUND,* which does not play a transforming role in our model (Figure S5J,K). Therefore, our study affirms the role of *FOSL1*, and uncovered *FOSL2*, *FOS* and *JUNB* as key members of the AP-1 network that mediates the malignant transformation of PDAC. While recent patient sample scRNA-seq nominates *FOSL2* as highly upregulated and a prognosis marker of PDAC^39^, our PO platform complements the patient sequencing data by elucidating the role of *FOSL2* in driving the malignant features associated with PDAC transformation and delineates the differing activities of AP-1 factors that have been viewed as functionally redundant in the context of human PDAC.

### *TET1* loss coincides with the suppression of pancreatic lineage program

Motifs related to TET1 and its binding partners, EGR1 and MECP2^40–42^, were consistently enriched in genomic regions losing accessibility in initial transformation in *KCP* POs, and regions that further lose accessibility during the progression of PDAC in motif analysis comparing *KCPS* and *KCP* POs (Figure 3K,L, S4O,P). Previous studies by our group and others have established the function of TET proteins and their partners in safeguarding normal pancreatic development programs^43,44^. In *KCP* and *KCPS* POs, we observed transcriptional downregulation of *TET1* but not *TET2* or *TET3* (Figure 5A). In mouse PanIN and PDAC organoids, a comparable finding was observed (Figure 5B, S6A). To investigate the clinical relevance of this finding, we analyzed a published single-cell RNA-seq dataset from 24 patient PDAC and 11 normal samples^45^. Using the expression pattern of either normal or malignant epithelial markers, we identified epithelial populations (Figure S6B-E), re-clustered these cells to define normal, abnormal, and malignant epithelia representing the trajectory of PDAC development based on the expression of malignant marker *MUC1*, and validated the approach by the expression pattern of additional malignant markers (*FXYD3*, *KRT7, KRT19* and *SLPI*) and normal epithelial markers (*CFTR*, *AMBP*, *FXYD2* and *HNF1B*) in the subclusters (Figure 5C-E, S6F,G)^45^. We confirmed the decrease of *TET1* (but not *TET2* or *TET3)* expression in malignant epithelia, while expression of pancreatic lineage TFs *ONECUT1* and *PDX1* was also markedly reduced (Figure 5F-J, S6H).

**Figure 5.**
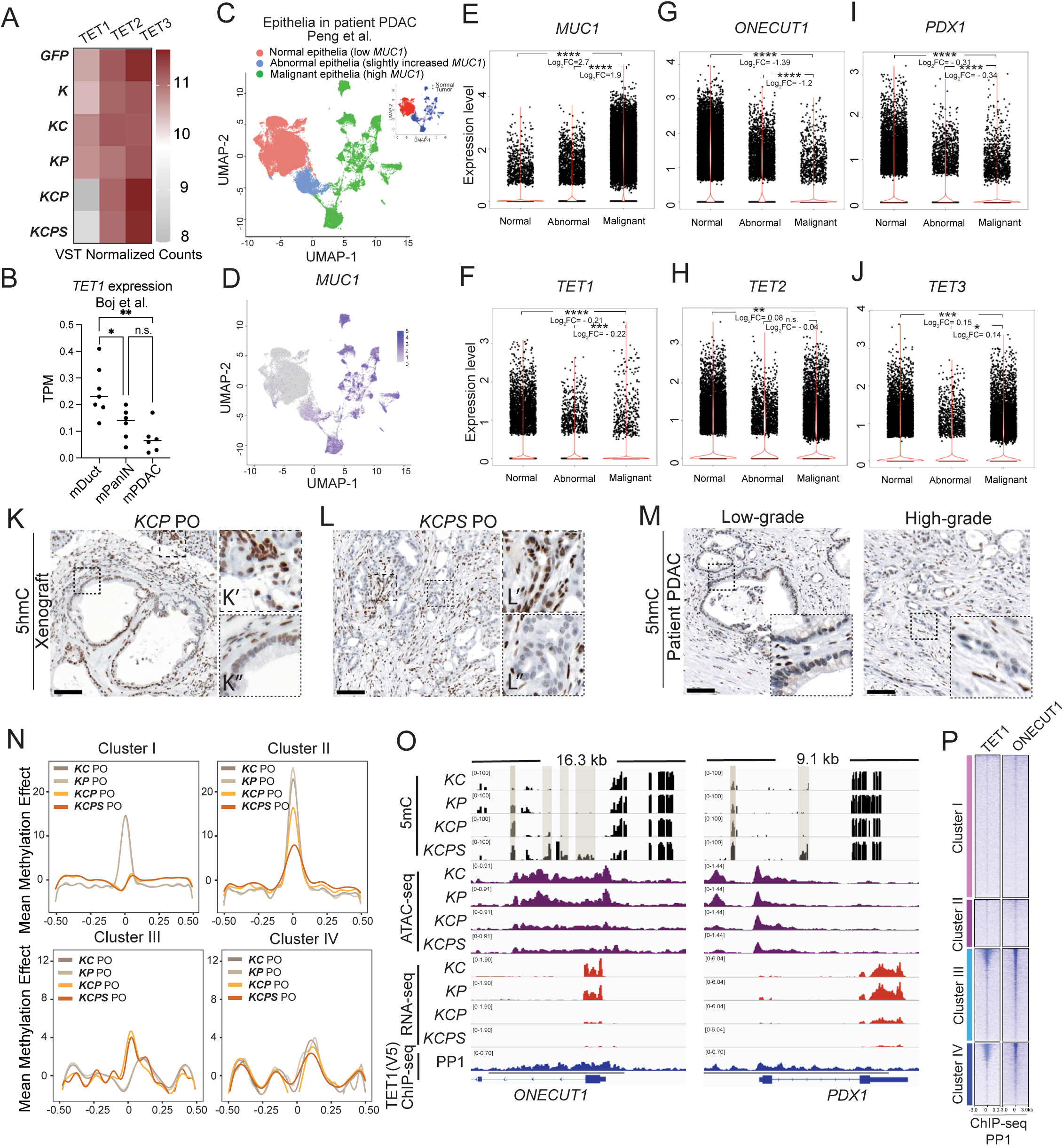
*TET1* loss coincides with the suppression of pancreatic lineage program. (A) A heap map of normalized counts of *TET1*, *TET2* and *TET3* mRNA in POs of indicated genotype was presented. (B) Expression level of *TET1* in mouse normal duct, PanIN and PDAC organoids from Boj et al. was displayed, one-way ANOVA, *Padj<0.05, **Padj<0.01. (C-J) Epithelial cells identified in patient PDAC and normal samples were re-clustered to reveal three major clusters indicating the trajectory of PDAC development based on the expression of multiple malignant epithelial markers including *MUC1*: normal (*MUC1* low), abnormal (*MUC1* slightly increased) and malignant (*MUC1* high) (C-E). Fold change and p values of gene expression in the three clusters was determined using Model-based Analysis of Single-cell Transcriptomics (MAST) package. The p values were calculated by Wald tests after maximum likelihood estimation (E-J). * Padj<1e-5, ** Padj <1e-50, *** Padj <1e-100, **** Padj <1e-200. (K-M) IHC against 5hmC was performed on *KCP* and *KCPS* xenografts (K,L), and low and high-grade patient PDAC samples (M). Normal ducts in top insets (K’,L’) served as positive control. Lower insets (K”, L”) showed staining details in the tumor epithelia. Representative of 3 tumors. Scale bars: 200μm. (N) EPIC-methyl capture sequencing was performed on *KC*, *KP*, *KCP* and *KCPS* POs, n=2 per genotype. DNA methylation signals associated with each cluster defined in Figure 3C were plotted. (O) Genomic visualization of methylation signals was presented alongside ATAC-seq and RNA-seq tracks. Hypermethylated differential methylated regions (DMRs) in *KCP* and *KCPS* POs vs *KC* and *KP* POs and peaks of TET1 binding were indicated. (P) ChIP-seq were conducted against TET1 (V5) and ONECUT1 in PP1 cells. The binding signals were plotted based on the defined genomic regions in Figure 3C and reordered by the binding signal.

Conversion of 5-methylcytosine (5mC) to 5-hydroxymethylcytosine (5hmC) by TET family methylcytosine dioxygenases plays an important role in active DNA demethylation^46,47^. In *KCP* and *KCPS* xenograft tumors, IHC staining detected a decrease of 5hmC signal in tumor epithelia compared to normal ducts. Similarly, low and high-grade patient PDAC tumors exhibited only weak 5hmC signal in tumor nuclei (Figure 5K-M). This suggested that the loss of 5hmC represents an early event in PDAC transformation^48^, coinciding with the decrease of *ONECUT1* expression. Methyl Capture assays revealed a significant decrease of DNA methylation in Cluster I regions (OR*_KCP_*_vs*KC*_=0.735, OR*_KCP_*_vs*KP*_=0.806, *p*<0.0001), and a significant increase of methylation in Cluster III regions in *KCP* PO as compared to *KC* and *KP* POs (OR*_KCP_*_vs*KC*_ =1.538, OR*_KCP_*_vs*KP*_ =1.370, *p*<0.0001, Figure 5N). Specifically, regions linked to PDAC markers *MUC5AC*, *S100P*, and *MMP7* exhibited a decrease in methylation in *KCP* and *KCPS* POs, concomitant with the opening of their chromatin structure (Figure S6I). Hypermethylated regions were identified in loci associated with the essential pancreatic lineage TFs *ONECUT1* and *PDX1*, and the increase of methylation in these regions was accompanied by a progressive decrease in chromatin accessibility and mRNA expression during PDAC development in *KCP* and *KCPS* POs (Figure 5O). Together, these data support that DNA methylation is dynamic in PDAC development in association with both acquired and repressed cell fates.

To determine the specificity of TET1 activity, we conducted ChIP-seq in PP1 expressing V5-tagged TET1 and confirmed the enrichment of TET1 binding in Cluster III and IV regions, including the loci associated with *ONECUT1* and *PDX1* (Figure 5P). Notably, binding of TET1 spanned the major hypermethylated regions identified in *ONECUT1* and *PDX1* loci in *KCP* and *KCPS* POs (Figure 5O). Therefore, *TET1* likely plays a crucial role in the maintenance of the pancreas lineage program, which is suppressed during malignant transformation.

### ERK activates oncogenic AP-1 and suppresses TET1-mediated pancreatic lineage program in early PDAC transformation

MEK/ERK pathway is a key driver of PDAC and other endoderm cancers downstream of oncogenic *KRAS*^49,50^. To examine how the oncogenic KRAS signaling influences the transcriptome and epigenetic reprogramming during human PDAC tumorigenesis, we applied the ERK inhibitor Temuterkib to *KCP* POs^51^. Acute treatment with Temuterkib for 4 days did not cause immediate cell death but significantly reduced the protein levels of *FOS*, *FOSL2* and *JUNB* (Figure 6A, S7A-C). Globally, ERK inhibition partially reversed the malignant transcriptome of *KCP* POs (Figure 4C, 6B). Among the previously defined 2105 TFM UP genes, 571 were downregulated by ERK inhibition, including multiple PDAC markers, *MUC5AC*, *TFF1*, *ANAX10,* and *S100P,* validated by RT-qPCR. Conversely, 529 out of 2631 TFM DN genes exhibited significant upregulation, including *ONECUT1* and *TET1* (Figure 6B, S7D, Table S6).

**Figure 6.**
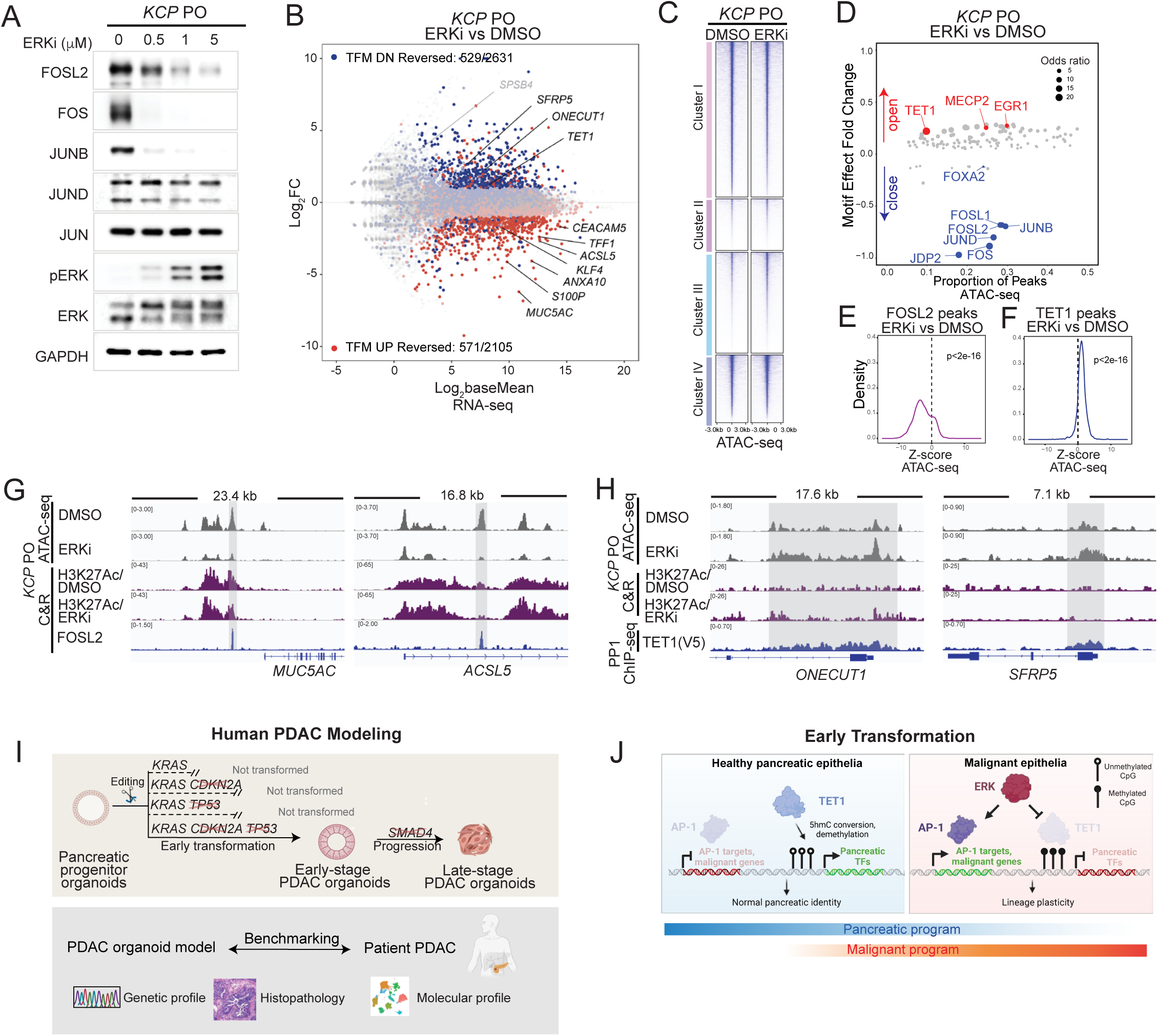
ERK activates oncogenic AP-1 and suppresses *TET1*-mediated pancreatic lineage program in early PDAC transformation. (A) *KCP* PO were subjected to Western blot analysis after treatment with ERK inhibitor Temuterkib for 4 days. (B) RNA was extracted from *KCP* POs treated as in (A) for RNA-seq. Differentially expressed gene analysis was conducted as in Figure 4C-H. (C) A tornado plot of ATAC-seq signal of *KCP* PO treated with Temuterkib or DMSO was displayed in the context of the genomic clusters identified in Figure 3C. (D) Motif analysis comparing Temuterkib vs DMSO-treated *KCP* PO was performed as in Figure 3K,L. (E,F) Density plots were generated to visualize the ATAC-seq signal ratio in Temuterkib and DMSO-treated *KCP* POs in FOSL2 and TET1 bound regions across the entire genome. Wilcoxon signed rank test was performed. (G,H) Genomic visualization of ATAC-seq signals in loci bound by FOSL2 (G) or TET1 (H) in *KCP* POs treated with Temuterkib or DMSO. Combined ATAC-seq, H3K27ac and FOSL2 C&R (G) or TET1 ChIP-seq in PP1 (H) tracks were presented. Statistically called FOSL2 and TET1 bound regions (IDR<0.01) were highlighted. (I,J) Schematic model depicted PDAC neoplastic transformation and progression driven by cumulative genetic mutations, as validated by histopathology of xenografts and multi-omics profiling of POs in culture (I), and the epigenetic reprogramming by oncogenic AP-1 and the loss of TET1-mediated lineage barrier during PDAC transformation downstream of ERK signal (J).

When examining the chromatin accessibility landscape upon treatment with Temuterkib in *KCP* POs, we observed a significant loss of accessibility in Cluster I (transformation open) regions, and re-opening of Cluster III (transformation closed) regions (Wilcoxon signed rank test p<2x10^-16^, Figure 6C, S7E,F). Motif analysis comparing *KCP* POs treated with ERK inhibitor or DMSO indicated chromatin regions gaining and losing accessibility were enriched with motifs of TET1 and AP-1 factors, respectively (Figure 6D), demonstrating that ERK inhibition impacts chromatin not only downstream of AP-1 but also associated with TET1. To further establish the role of oncogenic AP-1, TET1 and ONECUT1 in ERK-driven transformation of *KCP* POs, we analyzed the chromatin accessibility associated with regions bound by FOSL2, FOS and JUNB in *KCP* POs (Figure 4I), and TET1 or ONECUT1-bound regions identified in PP1 (Figure 5P). Genome-wide, a significant decrease of chromatin accessibility in the oncogenic AP-1-bound regions and gain of chromatin accessibility in TET1 and ONECUT1 bound regions were observed (Wilcoxon signed rank test p<2x10^-16^ in all cases, Figure 6E,F, S7I-K), consistent with their expression pattern upon ERK inhibition in *KCP* POs. Furthermore, chromatin accessibility of oncogenic AP-1-bound regions exhibited a more drastic decrease compared to unbound regions in Cluster I (Figure S7L-N). Similarly, gains of accessibility were more prominent in TET1 bound versus unbound regions in Cluster III, while ONECUT1 bound regions did not exhibit preferential opening in Cluster III regions (Figure S7O,P). ERK inhibition repressed Cluster I regions associated with *MUC5AC* and *ACSL5* which correlated with binding of oncogenic AP-1 factor *FOSL2* in *KCP* POs (Figure 6G). Conversely, TET1-bound Cluster III regions associated with *ONECUT1* and *SFRP5* were reactivated by ERK inhibition (Figure 6H). These findings indicated that ERK signaling not only mediates the activation of oncogenic AP-1 factors but enforces repression of *TET1* and thus the pancreas lineage program in *KCP* POs. Together, these two processes function in concert to drive the cellular plasticity during malignant transformation of human PDAC.

## Discussion

Leveraging hPSC-derived pancreatic progenitor organoids, we modeled the neoplastic transformation and progression of PDAC by combinatorial introduction of driver genetic events. Specifically, the combination of *KRAS*, *CDKN2A*, and *TP53* mutations, induced cellular transformation and resulted in PanIN and moderately differentiated PDAC upon orthotopic transplantation. Subsequent deletion of *SMAD4* led to poorly differentiated PDAC *in vivo*, consistent with the correlation between *SMAD4* loss and the emergence of high-grade PDAC histology and advanced squamous-like subtype in molecular profiling^11,48^. Mechanistic investigation revealed two crucial processes governing cellular plasticity during early malignant transformation: the activation of oncogenic AP-1 factors and epigenetic reprogramming of regions associated with trans-lineage/malignant features, and the suppression of a TET1-mediated pancreatic lineage program, two integral aspects of plasticity regulated by ERK signaling in PDAC (Figure 6I,J).

Our human PO model shows that the presence of either intact *CDKN2A* or *TP53* constitutes a barrier to PDAC transformation in oncogenic *KRAS* expressing human pancreatic progenitors. To date, no mouse model systems have established the requirement for *CDKN2A* loss in addition to *KRAS* and *TP53* alterations for PDAC development, although the *p16/CDKN2A* protein product is consistently lost in human PDAC, and the combination of *CDKN2A* and *TP53* mutations occurs in 68%∼78% of all patient PDAC samples^9,52^, indicating the non-redundant roles of the two TSGs in human. Similarly, multiple TSG deletions have been introduced to achieve complete cellular transformation in modeling of melanoma in human cells compared to their mouse counterparts^53,54^. This observation suggests human cells are inherently more refractory to oncogenic signals, though more cancer models need to be examined to access the generalizability of this phenomenon. Additionally, while no further genetic events are reported to be required when well-differentiated mouse tumors progress to poorly-differentiated PDAC upon the tumorigenesis^25,26^, progression to high-grade PDAC in our model requires genetic alterations, such as *SMAD4*, a finding that phenocopies human disease^11,48^, and highlights a limitation across the routinely used autochthonous mouse models in recapitulating the driver genetic events in PDAC initiation and progression.

It is noteworthy that even in the absence of exposure to the tumor microenvironment, *in vitro* cultured *KCP* and *KCPS* POs could recapitulate the transcriptomic and epigenetic signature of PDAC. This emphasizes the significance of cell-autonomous changes in malignant transformation and progression, at least in our organoid-based model. A pancreatitis-induced inflammation program is linked to susceptibility to oncogenic transformation in pancreatic epithelia^15^. Molecules involved in the interplay between tumor microenvironment and epithelial populations, as well as cell-intrinsic TFs and distinct patterns of epigenetic remodeling have been reported to participate in this process^16,18,55,56^. Our finding that pancreatic progenitor organoids faithfully reproduce the oncogenic transformation under clinically relevant genetic profiles^5,9^. suggests that signals induced by pancreatitis eventually converge on the onset of embryonic progenitor-like program in the epithelia, which confers conduciveness to neoplastic transformation. To date, multiple attempts have been launched to recapitulate PDAC tumorigenesis in human pancreatic epithelia by reconstituting the precursor or PDAC mutation profiles^57–60^. These efforts have resulted in the formation of PanIN or PDAC in a variety of contexts. However, the permissiveness to oncogenic transformation in the epithelia, as well as the necessity of TSG alterations in PDAC initiation has been inconsistent across these systems, as “*KCPS”* mutations in mature pancreatic ductal cells are unable to induce PDAC^57^, while oncogenic *KRAS* alone gives rise to full spectrum of PanIN/PDAC in hPSC-derived ductal and acinar cells^59,60^. Compared to these studies, our model reproduces the human genetic requirements of early and advanced PDAC, likely due to the selection of pancreatic progenitors as cell of origin, which closely mimic the inflamed epithelia in pancreas.

Tumor progression and drug resistance in endoderm malignancy has been associated with cellular plasticity and targeting underlying mechanisms in distinct context has revealed novel vulnerabilities in cancer cells^61,62^. In contrast to the focus on ‘acquired plasticity’ in both tumor initiation and progression, the molecular mechanisms underlying the repression of original cell identity (i.e. loss of lineage specification) initiated during early PDAC transformation and continued throughout disease progression remain understudied. Loss of 5hmC has been observed in both pre-neoplastic and PDAC lesions, indicating impaired DNA demethylation activity is an early marker in malignant transformation^48^. Our model fully recapitulates the suppression of 5hmC and hypermethylation of essential pancreas lineage TFs in early and late PDAC, and reveals that *TET1,* through safeguarding the lineage program, acts as an epigenetic barrier against tumorigenesis. The commitment to pancreatic lineage is silenced accompanying *TET1* loss upon PDAC onset. Although rarely mutated in PDAC, hypermethylation and repression of pancreatic lineage TFs are linked to tumor progression and poor patient prognosis^28,48,63,64^. Notably, TFs significantly upregulated upon pancreatic lineage specification in normal development, including *ONECUT1*, *PDX1* and *NR5A2,* exhibit decreased expression in transformed POs and cancerous epithelia in clinical samples^45^. The downregulation of these TFs is potentially selected during tumor evolution, as they have been shown or proposed to possess tumor suppressor functions in neoplastic transformation^34,35,65,66^, while restoration of these pancreatic TFs and normal cell differentiation in tumor cells could provide therapeutic benefits as reported in the treatment of leukemia^67^. Current efforts to develop targeted therapies for PDAC primarily focus on mutations and alterations in established tumors. However, prevention and early intervention strategies remain critically lacking for individuals with high-risk profiles. Our findings reveal the interplay between oncogenic signaling and lineage restriction pathways in tumorigenesis, and support TET activation and lineage restoration as promising early intervention strategies for those at high risk of developing PDAC, offering a promising avenue to improve outcomes for this devastating disease.

## Data and code availability

Analysis was performed with published tools as described in the Methods. Scripts for generating computational results are available upon request. The original sequencing data is available from Gene Expression Omnibus under accession number GSE273073, which will be released upon publication.

## Acknowledgements

We acknowledge the use of the Molecular Cytology Core, Integrated Genomics Operation Core and the Antibody & Bioresource Core of Memorial Sloan Kettering Cancer Center. We thank the David M. Rubenstein Center for Pancreatic Cancer Research and the Society of MSK for support. We thank Nicolas Lecomte, Xianlu Laura Peng and Jen Jen Yeh for assisting with additional experiments not included in the manuscript. This work is partially supported by grants to D.H. from NIH (R01DK096239, R01HD111256), the Starr Tri-I Stem Cell Initiative (2016-032), Geoffrey Beene Cancer Research Center, Shipley Foundation, Cycle for Survival Funds from the David M. Rubenstein Center for Pancreatic Cancer Research, and MSKCC Special Projects Committee. Additional supports include a MSKCC Cancer Center Support Grant from the NIH (P30CA008748); a Geoffrey Beene Cancer Research grant (to A. Hall), a Tri-Institutional Starr Stem Cell Scholars Fellowship (to X.Z.), a postdoctoral fellowship from a NYSTEM training grant from the Center for Stem Cell Biology of the Sloan Kettering Institute (DOH01-TRAIN3-2015-2016-00006, to D.Y.), a Beatrice P. K. Palestin Fellowship and a Bruce Charles Forbes Fellowship (to D.L.); and a National Institutes of Health T32 training grant T32GM008539 (to S.J.K.).

## Author contributions

X.Z. initiated the project under the guidance of Alan Hall. F.G., X.Z., and J.P. constructed plasmids. X.Z. performed the hPSC gene-editing, cell differentiation, PO culture, Western and Southern blot, immunofluorescence. X.Z., R.L. and F.C.P. performed orthotopic transplantation. X.Z., J.R.D. and R.W.R. performed mouse husbandry. X.Z., R.K. and J.R.D. performed immunohistochemistry. S.U., G.A. and A.Y. reviewed the histology of xenografts. S.U. quantified AP-1 immunohistochemistry on MSKCC tumor microarray. V.G.A. identified human PDAC samples, sorted nuclei, and prepared ATAC-seq libraries; A.G. prepared and analyzed human ATAC-seq libraries. S.D.L. provided funding and oversight of studies involving intra-pancreatic injections and ATAC-seq analysis of PDAC tumors. X.Z. and G.D. generated AP-1 sgRNA virus and X.Z. performed perturbation screen in POs. X.Z., D.Y. and D.L. performed ChIP-seq. W.W.and H.S.C. performed the data analysis on bulk RNA-seq, ATAC-seq and ChIP-seq. W.W., S.J.K. and D.T. analyzed RNA-seq and performed GSEA analysis. W.W. analyzed TCGA and GTEx data, and R.L. analyzed patient scRNA-seq data. H.P. analyzed EPIC methyl capture data. R.C., C.L., and D.H. co-supervised the study. X.Z. and D.H. wrote the manuscript. All authors reviewed and edited the final manuscript.

## Supplemental information

Document1 Supplemental Figures 1-7 with figure legends

Table S1 Histology summary of xenograft assay

Table S2 Summary of meta-analysis and top up and down-regulated genes in *KCP* vs *GFP* POs used for GSEA (related to Figure 2)

Table S3 Summary of clustering of top 3000 variable genes across all genotypes of POs (related to Figure 2)

Table S4 Summary of patient PDAC samples used for ATAC-seq (related to Figure 3)

Table S5 Summary of top variable ATAC-seq peaks in all genotypes of POs (related to Figure 3)

Table S6 Summary of differentially expressed gene analysis in AP-1 knockout and ERKi treated *KCP* POs (related to Figure 4,6)

Table S7 List of sgRNA target sequences, primer sequences for genotyping and RT-qPCR Table S8 List of primary antibodies

## Methods

### Generation of constructs

To create the DNA template for TALEN-mediated gene-editing at the first intron of PPP1R12C at the AAVS1 locus, HA-L-SA-2A-Puro^R^-GFP-TRE-HA-R and HA-L-SA-2A-Puro^R^-GFP-KRAS^G12V^-TRE-HA-R donor constructs were generated by cloning the coding sequence of GFP or GFP-KRAS^G12V^ from a lentiviral expression construct from Alan Hall Lab ^68^ into pDONR201 entry vector and then transferred into puro-iDEST vector (Addgene #75336)^69^ through Gateway cloning. HA-L-SA-2A-Neo^R^-Cag-M2rtTA-Cas9-TRE-HA-R constructs were generated by cloning coding sequence of Cas9 with 3xFLAG tag from a pAAVS1-PDi-CRISPRn construct (Addgene #73500, gift from Bruce Conklin lab)^70^ into the multiple cloning sites of a pAAVS1-NDi-MCS vector under TRE3g promotor (gift from Bruce Conklin lab). Neo^R^ resistance cassettes was under 2A sequence, and rtTA3G cassette were under CAG promotor in the same construct.

To deliver the *CDKN2A* and *TP53* sgRNAs into the hPSCs, pWPXL-Hygro (DEST) vector was constructed from pWPXL lentiviral backbone (Addgene #12257). In summary, the eGFP CDS was removed by PmeI-SpeI digestion and replaced with a PmeI-attR1-CamR-ccdB-attR2-SpeI Gateway destination cassette PCR amplified on pDEST-17 Vector (Invitrogen). The plasmid was linearized again using PacI digestion and a PacI-Em7-Hygro-SV40pA-PacI cassette was PCR amplified on Hygro-Cas9 donor (Addgene #86883) and inserted at this site. pPGKenCh-sgRNA (ENTRY) vector was constructed from piCRg Entry (Addgene #58904) by partial PCR amplifying and self-ligating to remove the NLS-Cas9-bGHpA insert thus creating an attL1-U6-BbsI-tracr-attL2 Gateway entry cassette. *CDKN2A* or *TP53* sgRNAs were cloned into this entry vector as described in Zhang Lab General Cloning Protocol for pX330 vector ^71^. The U6-sgRNA entry expression cassettes from pPGKenCh-*CDKN2A*Cr or pPGKenCh-*TP53*Cr were transferred into pWPXL-Hygro(DEST) using LR clonase (Invitrogen) to create iKO-*CDKN2A*Cr and iKO-*TP53*Cr sgRNA expression plasmids. iKO-*CDKN2A*Cr was digested with AscI, 5’ overhangs were removed with Mung Bean Nuclease and the PCR amplified U6-*TP53*Cr-tracr cassette from iKO-*TP53*Cr vector was cloned downstream of the U6-*CDKN2A*Cr expression cassette to create iKO-*CDKN2A*Cr-*TP53*Cr sgRNA expression plasmid.

To deliver AP-1 specific sgRNA into *KCP* POs, a lentiGuide-Blast sgRNA expression vector was constructed by replacing the puromycin resistance cassette of lentiGuide-Puro vector (Addgene #52963) with a Blasticidin resistance cassette from lenti-dCAS9-VP64-Blast vector (Addgene #61425) by In-Fusion® Cloning (Takara Bio, 638948). AP-1 sgRNA sequences were cloned into the vector following the sgRNA cloning protocol ^71^. A WPXL-Luc2tdT was a gift from Wenjun Guo lab ^72^.

### *In vitro* transcription of sgRNAs

To generate *SMAD4* sgRNA *in vitro*, a T7 promoter was added to sgRNA templates by PCR amplification on piCRg Entry vectors (Addgene #58904) using CRISPR-specific forward primers and a universal reverse primer gRNA-R (5’-AAAAGCACCGACTCGGTGCC-3’). T7-sgRNA PCR products were used as templates for *in vitro* transcription using the MEGAshortscript T7 kit (Life Technologies, AM1354). The resulting sgRNAs were purified using the MEGAclear kit (Life Technologies, AM1908), eluted in RNase-free water and stored at -80°C until use.

### Tissue culture

HUES8 hPSCs (NIHhESC-09-0021) were cultured on irradiated mouse embryonic fibroblasts (iMEFs) feeder layers in DMEM/F12 medium (Life Technologies, 11320) supplemented with 20% KnockOut Serum Replacement (Life Technologies, 10828028), 1X MEM Non-Essential Amino Acids (Life Technologies, 11140), 1X GlutaMAX (Life Technologies, 35050079), 100U/ml Penicillin and 100 μg/ml Streptomycin (Gemini, 400-109), 0.055 mM 2-mercaptoethanol (Life Technologies, 21985023) and 10 ng/ml recombinant human basic FGF (EMD Millipore, GF003AF-MG). Cells were maintained at 37 °C with 5% CO2 and passaged at 1:12 split ratio every 4 - 5 days using ReLeSR (Stem Cell Technologies, 05872). 5 μM Rho-associated protein kinase (ROCK) inhibitor Y-27632 (Selleck Chemicals, S1049) was added into the culture medium when passaging cells. H1 hPSCs (NIHhESC-10-0043) were maintained in the chemically defined feeder-free, serum free Essential 8 (E8) medium (Thermo Fisher Scientific, A1517001) on vitronectin (Thermo Fisher Scientific, A14700) coated plates. Cells were passaged every 4 days at 1:20∼25 ratio using 0.5 mM EDTA (KD Medical, RGE-3130) to dissociate cells. 293T cells used for lentivirus production were maintained in DMEM supplemented with 15% fetal bovine serum, 1X GlutaMAX, 1X MEM NEAA and 1mM Sodium Pyruvate (Life Technologies, S8636).

### Generation of hPSC lines that enable doxycycline inducible mutations associated with **PDAC**

2mg each of AAVS1-TALEN-L and AAVS1 TALEN-R constructs targeting the first intron of PPP1R12C at the AAVS1 locus ^73^ were co-electroporated with DNA template (a) HA-L-SA-2A-Puro^R^-GFP-TRE-HA-R or HA-L-SA-2A-Puro^R^-GFP-KRAS^G12V^-TRE-HA-R and template (b) HA-L-SA-2A-Neo^R^-Cag-M2rtTA-Cas9-TRE-HA-R by AMAXA nucleofector (Lonza). The two TALEN constructs create site specific DNA cleavage, allowing the insertion of DNA expression cassettes into both alleles of the human AAVS1 site through homology-directed repair. Following TALEN based insertion of GFP-*KRAS^G12V^*/ *Cas9* or *GFP/Cas9* expression cassettes, genomic DNA from cloned hPSCs underwent SphI restriction enzyme digestion overnight, followed by Southern blot analysis. An internal probe recognizing 5’ homology arm (HA-L) was designed to detect fragments of proper size (6492bp DNA fragment for wild-type genome, 3781bp for *GFP* or *GFP*-*KRAS^G12V^*cassettes and 3519bp fragments for *Cas9* cassette). Clones exhibiting additional fragments, indicating random insertion, were excluded from subsequent experiments.

hPSC clones verified by Southern blot were treated with doxycycline and validated by Western blot to confirm the expression of *GFP*/*GFP-KRAS^G12V^*and *Cas9*. Validated *KRAS*^G12V^/*Cas9* hPSC lines were then transduced with lentivirus expressing sgRNAs targeting *CDKN2A*, *TP53* or both genes in tandem. T7E1 assays were performed to select hPSC clones with high knockout efficiency for pancreatic differentiation and organoids production.

### Directed pancreatic differentiation

NKX6.1+ pancreatic progenitors (PP2) were produced following stepwise specification of 1) definitive endoderm (DE), 2) primitive foregut (FG), 3) early PDX1^+^pancreatic progenitor (or posterior foregut, PP1) and 4) NKX6.1^+^ pancreatic progenitor (or pancreatic endoderm, PP2) from hPSCs. For differentiation of genome edited hPSC for pancreatic progenitor organoid development, hPSCs were first adapted to mTeSR medium (Stem Cell Technologies, 85850) for two weeks in Matrigel (Corning, 354230) coated culture dish. One day before the differentiation, hPSCs were digested into single cells using Accutase (Stem Cell Technologies, 07920) and plated on Matrigel coated dish to reach a confluence of 80% on the day of differentiation. STEMdiff™ Pancreatic Progenitor Kit (Stem Cell Technologies, 05120) was employed following the instructions from the manufacturer’s instruction except that one more day of medium 1B treatment was applied in stage 1 before the specification of DE, which greatly enhanced the PP2 differentiation efficiency as monitored by percentage of PDX1/NKX6.1 positive cells.

Large scale PP2 differentiation for ChIP-seq was performed following the protocol previously described ^74^. Wild-type and TET1-V5 tagged H1 hPSC cells were maintained in E8 medium (Thermo Fisher Scientific, A1517001) in vitronectin (Thermo Fisher Scientific, A14700) coated dishes. The day before differentiation, cells were replated and maintained in E8 medium for one day to reach ∼80% confluence before the start of differentiation. Differentiation medium was prepared as previously described ^74^: S1/2 media: MCDB 131 medium (Life Technologies, 10372-019) supplemented with 1.5 g/l sodium bicarbonate (Fisher Scientific, s22060), 1× Glutamax (Thermo Fisher Scientific, 35050061), 10 mM Glucose (Sigma-Aldrich, G8769), 0.5% BSA (LAMPIRE Biological Products, 7500804). S3/4 media: MCDB 131 medium supplemented with 2.5 g/l sodium bicarbonate, 1× Glutamax, 10 mM Glucose, 2% BSA. hPSCs were rinsed with 1X DPBS (without Mg2+ and Ca2+) and first differentiated into DE using S1/2 media supplemented with 100 ng/ml Activin A (Bon Opus Biosciences) for three days and 5 mM CHIR99021 (Stemgent, 04-0004-10) for the first day. DE cells were rinsed with 1X DPBS (without Mg2+ and Ca2+) and then exposed to S1/2 media supplemented with 50 ng/ml of KGF (FGF7) (PeproTech, 100-19) and 0.25mM vitamin C (Sigma-Aldrich, A4544) for 2 days to reach GT stage. For differentiation toward PP1 stage, cells were exposed to S3/4 media supplemented with 50 ng/ml of FGF7, 0.25 mM vitamin C, 2µM retinoic acid (RA, Sigma-Aldrich, R2625), 200 nM LDN (Stemgent, 04-0019), 0.25 µM SANT-1 (Sigma, S4572), 200 nM TPB (EMD Millipore, 565740), and 1:200 ITS-X (Life Technologies 51500056) for two days. The cells were then further differentiated toward PP2 stage using S3/4 media supplemented with 50 ng/ml of FGF7, 0.25 mM vitamin C, 0.1 µM RA, 200 nM LDN, 0.25 µM SANT-1, 200 nM TPB, and 1:200 ITS-X for 4 days.

### Establishment of wild-type and *GFP*, *K*, *KC*, *KP*, *KCP* and *KCPS* PO

*In vitro* differentiated PP2 cells were dissociated in Accutase for 5min to achieve 3∼5 cell clusters, washed once in ice cold DMEM-F12 base medium (Life Technologies, 11330057) and resuspended in growth factor reduced Matrigel (Corning, 354230). A 30 ml Matrigel dome was applied to the bottom of each well of non-tissue culture treated 48 well plate (Thermo Fisher Scientific,150787) and kept in 37°C incubator for 10 min for Matrigel to solidify. DMEM-F12 base medium was supplied with Glutamax, Penicillin/Streptomycin, B27^TM^ supplement (Life Technologies, 17504044), 1mM N-acetyle-L-cystein (Sigma-Aldrich, A9165-25G), 10mM Nicotinamide (Sigma-Aldrich, N0636-100G), 0.5mM A83-01 (Tocris, 2939), 100ng/ml hNoggin (Peprotech, 120-10c), 50ng/ml hEGF (R&D Systems, 236-EG-200), 10nM hGastrin I (Sigma-Aldrich, G9020-250UG), 100ng/ml hFGF10 (PeproTech,100-26), 500ng/ml Rsp1 (as gift from Albert Einstein College of Medicine). 0.4ml of medium was applied to each well. 5 μM Y-27632 was present in the medium during the first 4 days of culture after passaging, which was replaced with fresh medium without Y-27632 (Selleck Chemicals, s1049) at the end of the 4^th^ day. Organoids were split every 7 days. A distinct population of mesenchyme-like cells, originating from non-specified cells during pancreatic endoderm differentiation was observed between week 4 and 8 and manually removed from the culture.

To induce the PDAC associated mutation in POs, PP2 cells differentiated from previously genome edited hPSC (*GFP*, *K*, *KC*, *KP*, *KCP*) were adapted to the Matrigel culture for 1 week followed by 3mg/ml doxycycline (Sigma-Aldrich, D3072) treatment in the 2^nd^ week, 1mg/ml doxycycline in the 3^rd^ week and 0.5mg/ml doxycycline thereafter for *GFP*, *K, KC* and *KP* POs. *KCP* POs were maintained in 0.05 mg/ml doxycycline. To establish *KCPS* POs, CRISPR sequencing validated *KCP* POs were dissociated by Accutase into single cells and 300ng of *in vitro* transcribed *SMAD4* sgRNA was electroporated (Lonza, Amaxa) into 0.5 million dissociated *KCP* PO cells. 10ng/ml of TGF-b was applied in the culture medium to select for *SMAD4* knockout POs for 2 weeks or until the non-electroporated POs all died. *SMAD4* knockout efficiency was verified by CRISPR sequencing in the surviving *KCPS* POs. *KCPS* POs were maintained in 0.1 mg/ml doxycycline.

### Multi-lineage differentiation of PO

Multi-lineage differentiation of POs towards acinar, ductal and endocrine cells were conducted based on a modified protocol from Trott et al ^24^. DMEM-F12 medium was supplied with 2.5 g/30 mL BSA, Glutamax, Penicillin/Streptomycin, B27^TM^ supplement to constitute the base medium for multi-lineage differentiation. PO cultured in Matrigel for 8∼12 weeks were split and re-plated in PO medium for one day, washed two times with base medium for differentiation, and treated with 4.5 mM RA, 1.5 mM DAPT (Sigma, D5942) and 100 mM BNZ PKA activator (Sigma, B4560) for the first 4 days. On day 4 and day 7, base medium containing 4.5 mM RA and 0.375 µM SANT-1 was applied. POs were harvested and RNA extracted for analysis on day 10.

### Immunofluorescence staining

*In vitro* differentiated PP2 cells were fixed in 4% paraformaldehyde diluted from 32% paraformaldehyde (Thermo Fisher Scientific, #50980495) for 10 minutes at room temperature and permeabilized with 0.1% Triton X-100 in PBS. After washing with PBS three times for 5 minutes each, cells were blocked in 5% BSA in PBS buffer for 30 minutes. 3D cultured POs were directly fixed in 4% paraformaldehyde for at least 30 minutes in room temperature. Permeabilization was performed in 1x IF solution (Nacl 1.3M, Na2HPO4 0.13M, NaH2PO4 0.03M, NaN3 0.5%, BSA 1%, Triton-X 100 2%, Tween-20 0.4%) for one hour, followed by one hour blocking in 5% BSA in PBS. Fixed cells were incubated with primary antibodies diluted in blocking solution overnight at 4°C. After three washes in PBS, secondary antibodies diluted in blocking solution were applied for 1hr, followed by DAPI staining for 15min at room temperature. Confocal images were taken on a Leica TCS SP5 inverted microscopy.

### Orthotopic transplantation

Intrapancreatic transplantation was performed under protocol approved by MSKCC IACUC. hPSC derived POs were isolated from the Matrigel using Cell Recovery Solution (Corning, 354253), washed with ice-cold PBS, physically sheared into pieces by triturating through fire-polished glass pipettes, and resuspended in 30 ml of Matrigel. 6 to 8 weeks old male NSG mice (Jackson Laboratories, 005557) were shaved and anesthetized using Isoflurane. An incision was made in the left abdominal side. Approximately 0.5 million organoid cells were injected into the tail region of the pancreas using insulin syringes (29 Gauge). The abdominal wall was sutured with absorbable Vicryl suture (Ethicon, J392), and the skin was closed with wound clips (AutoClip® System, Item No. 12020-00). Mice were fed doxycycline water the day before surgery until euthanized at the indicated time.

### Tissue processing, section and immunohistochemistry

Pancreas or tumor samples from xenograft experiments were fixed in 4% paraformaldehyde overnight at 4°C with rocking. Fixed samples were washed twice in PBS, preserved in 70% ethanol and submitted to the Molecular Cytology Core of MSKCC for automated tissue processing. Processed samples were embedded in paraffin and subjected to manual section. Section was performed at 5 mm per slide. Paraffin embedded patient PDAC sections or tumor arrays were sectioned at 5 mm per slides. Additional pre-sectioned PDAC tumor arrays were purchased from US Biomax (PA721a). Before use, slides were placed on sides in a 56°C oven for 1 hour, deparaffinized in Histo-Clear II solution (National Diagnostics, HS-200) followed by rehydration in 100%, 95% and 70% ethanol and H2O.

Hematoxylin and Eosin staining was performed in the Molecular Cytology Core of MSKCC. Rehydrated slides were stained in Harris Hematoxylin with acetic acid (Poly Scientific, S212A8OZ), differentiated in 0.3% acid alcohol, blued in PBS and stained in alcoholic Eosin Y Solution (Sigma-Aldrich, 102439). After Eosin staining, slides were rinsed in 95% and 100% ethanol before mounted in Permount mounting media (Fisher Scientific, SP15100). Alcian blue staining was performed using an Alcian Blue Stain Kit (IHC World, IW3000) per manufacturer’s instruction. Rehydrated slides were stained in Alcian Blue for 30 min, rinsed and counterstained with fast red for 10 min at room temperature. Stained slides were then dehydrated and mounted in Permount mounting media.

For immunohistochemistry staining, antigen retrieval was performed by boiling slides in antigen retrieval buffer (pH9, Thermo Scientific, 00-4956-58) for 20 min at 98 °C in microwave. Endogenous enzyme activity was blocked by dual enzyme block solution in the EnVision®+ Dual Link System-HRP kit (Agilent, K406511-2). Mouse and rabbit primary antibody was diluted in Antibody Diluent (Agilent, S0809) and slides were incubated in primary antibodies at 4 °C overnight. Anti- mouse and rabbit secondary antibodies conjugated with HRP-polymer (EnVision®+ Dual Link kit) were applied at room temperature for 45 min before visualization with substrate-chromogen solutions (EnVision®+ Dual Link kit). For staining of goat primary antibody (MMP7), 2.5% horse serum was applied for blocking purpose after dual enzyme block for 30 min at room temperature. Slides were incubated with primary antibody at 4 °C overnight. ImmPRESS^TM^ horse anti-Goat IgG HRP Polymer (Fisher Scientific, NC0461902) was applied for 45 min at room temperature before visualization with substrate-chromogen solutions (EnVision®+ Dual Link kit). All slides were counterstained in Harris Hematoxylin, dehydrated and mounted in Permount mounting media.

Automated immunohistochemistry detection of GFP and E-cadherin was performed on a Discovery XT processor (Ventana Medical Systems, Roche - AZ) in the Molecular Cytology Core of MSKCC. A mouse monoclonal anti-E-Cadherin antibody (BD Bioscience, cat. #610181) was applied as the primary antibody, followed by rabbit anti-mouse linker antibody (abcam, ab133469), biotinylated goat anti-rabbit IgG antibody (Vector labs, PK6101), and Streptavidin-HRP incubation. Primary chicken polyclonal anti-GFP antibody (Abcam, cat#ab13970) staining was followed by biotinylated goat anti-chicken IgG (Vector labs, T1008) and Streptavidin-HRP incubation. DAB detection kit (Ventana Medical Systems, Roche) was used to visualize the staining according to the manufacturer instructions. Slides were counterstained with hematoxylin and mounted in Permount.

### Flow cytometry

*In vitro* differentiated PP2 cells were dissociated using Accutase and resuspended in FACS buffer (5% FBS in PBS). Organoids were separated from Matrigel using the Cell Recovery Solution according to the manufacturer’s instruction and then dissociated into single cells by Accutase treatment. Single organoid cells were resuspended in FACS buffer before staining. LIVE-DEAD Fixable Violet Dead Cell Stain (Invitrogen, L34955) was used to label dead cells and staining was performed for 15 min at room temperature in FACS buffer. Fixation and intracellular staining were performed with Foxp3 Staining Buffer Set (eBioscience, 00-5523-00) following the manufacturer’s instructions. Permeabilization/fixation was performed at room temperature for 1 hour or 4 °C overnight. Primary and secondary antibody staining was performed at room temperature for 30 min in 1x permeabilization buffer. Cells were then analyzed using BD LSRFortessa or BD LSRII. Analysis was conducted using FlowJo v10 software.

### Southern blot

Southern blot was performed as previously described ^75^. Sequence from 5’- homology arm (HA-L) of both donors (HA-L-SA-2A-Puro^R^-GFP-KRAS^G12V^-TRE-HA-R and HA-L-SA-2A-Neo^R^- Cag-M2rtTA-Cas9-TRE-HA-R) was amplified by PCR to generate the internal probe using the PCR DIG Probe Synthesis Kit (Roche, 11636090910, forward primer 5’-AGGTTCCGTCTTCCTCCACT, reverse primer 5’-GTCCAGGCAAAGAAAGCAAG). 10mg of genomic DNA for each sample was digested with 20 U of SphI at 37 °C overnight, run on 1% TAE agarose gel, denatured, neutralized, and transferred overnight by capillarity on Hybond-N membranes (GE Healthcare, RPN203N) using 10x SSC transfer buffer. Hybridization with the internal probe was carried out overnight at 65 °C. Probes were detected by an AP-conjugated DIG-Antibody using CDP-Star (Roche, 11636090910) as a substrate for chemiluminescence as per manufacturer’s instructions.

### Western blot

Organoids were harvested using Cell Recovery Solution, rinsed with ice old PBS and lysed using cell lysis buffer (Cell Signaling Technology, 9803) containing cOmplete™ Mini Protease Inhibitor Cocktail (Roche, #05892791001). Protein concentration of each sample was measured using BCA Protein Assay Kit (Thermo Scientific™ 23227) and adjusted accordingly. Lysate was then mixed with NuPAGE® LDS Sample Buffer (4X, NP0007) and Sample Reducing Agent (10X, NP0009) and boiled for 5 min, before being loaded onto a NuPAGE™ Bis-Tris 4-12% protein gel (Fisher Scientific, NP0336BOX) and transferred to Immobilon®-P PVDF Membrane (EMD Millipore, IPVH00010). Membranes were blocked with 5% milk in 1x TBS (Cedarlane Labs, 42020060-1) supplied with 0.1% Tween-20 for 1 hour at room temperature. The membrane was incubated with primary antibodies overnight at 4°C, followed by incubation with HRP conjugated secondary antibodies at room temperature for 1 hour. ECL western blotting detection reagent (Amersham, RPN2236) was used to visualize the protein bands on HyBlot CL® Autoradiography Film (Thomas Scientific #1159T41) or directly on Amersham Imagequant 800 Western blot imaging system.

### RNA isolation and RT-qPCR

Organoids were lysed in 1 ml of TRIzol™ Reagent (Thermo Fisher Scientific, 15596-018) followed by application of 0.2ml of chloroform (Fisher BioReagents, BP1145-1) that separated RNA from DNA and protein. RNA from the aqueous solution was precipitated with 0.5ml of isopropanol, washed twice with 75% ethanol and re-suspended with DEPC water. For cDNA synthesis for quantitative real-time PCR, 1ug of RNA per sample was used per instruction from the manufacturer of a High-Capacity cDNA Reverse Transcription Kit (Thermo Fisher Scientific, 4368814). Equivalent of 10ng/4ng of total RNA was loaded in a 25 ml/10 ml qPCR reaction using the PowerUp SYBR Green Master Mix (Thermo Fisher Scientific, A25742) and run in a 96 well/384 well plate. Relative expression of target gene in an experimental sample versus the reference sample was calculated as: 2^-Δ CT(exp sample-ref sample)^ value/ mean of 2^-Δ CT(exp sample-ref sample)^ value of two housekeeping genes. *GAPDH*, *UBC9* or *RPL13A* were used as housekeeping genes in these experiments.

### RNA-seq and transcriptome analysis of PO samples

RNA of POs was submitted to MSKCC Integrated Genomics Operation Core. After RiboGreen quantification and quality control by Agilent BioAnalyzer, 500 ng of total RNA underwent polyA selection and TruSeq library preparation according to instructions provided by Illumina (TruSeq Stranded mRNA LT Kit, catalog # RS-122-2102) and run on a HiSeq 4000 as PE50 or on a NovaSeq 6000 as PE100 using the HiSeq 3000/4000 SBS Kit or NovaSeq 6000 S4 Reagent Kit (200 Cycles) (Illumina). Fastq files were first checked with FastQC (0.11.9). RNA-seq reads were quantified using Salmon (1.4.0) in hg19. Gene models from Ensembl’s GRCh37.p13 was used. After quantification, DESeq2 (1.30.1) was used for subsequent processing and identification of differentially expressed genes. A Wald Test was used to compare all pairwise conditions, and differentially expressed genes were reported at an adjusted p-value of 0.1.

### Principal component analysis of POs and clinical RNA-seq samples

Recount3 (1.0.7)^32^ was used to access processed counts for the TCGA-PAAD and GTEx V8 projects. Counts were downloaded and analyzed using gene models from Gencode v26 (G026). These pre-processed counts were used for further downstream analysis. The Monorail project, consisting of recount-pump (1.0.6) and recount-unify (1.0.4), was used to process the PO RNA-seq data according to the Recount3 specification. Then quantification was carried out using gene models from Gencode v26 (G026) for PCA projection. The PCA plot was generated by first performing PCA on the TCGA and GTEx samples using VST counts and then projecting the VST counts of the Recount3 processed PO data onto the first two principal components. To separate samples based on tumorigenicity of clinical samples, we first labeled samples according to their tumorigenicity and then used the PC1 and PC2 coordinates as input for a logistic regression model to define a separating hyperplane in the PC space by converting the coefficients of the logistic regression involving PC1 and PC2.

### POs expression heatmap and AP-1 knockout expression MA plot

To generate the RNA-seq heatmap of *GFP*, *K*, *KC*, *KP*, *KCP* and *KCPS* POs, the top 3000 variable genes were selected, and the VST counts were plotted with pheatmap package (1.0.12) The top 3000 variable genes were divided into 6 separate groups by unsupervised clustering into 2 sub-clusters over all the PO samples. Genes in the sub-cluster with overall up-regulated expression between *KCP* or *KCPS* and *GFP* POs were subdivided into 3 clusters, Cluster 1,2 or 3, based on if their log2 fold changes between *KCPS* and *KCP* were less than -2, -2 < 0 < 2, or >2. Similar approach was used to define Cluster 4,5,6 genes which displayed a general down-regulated expression between *KCP* or *KCPS* and *GFP* POs. To examine the effectiveness of AP-1 knockout at reversing AP-1 mediated gene expression changes, we compared the fold changes of the genes between *KCP* and *KP* to the fold changes of the genes between knockout and NT control experiments. The fold-changes were standardized by their standard errors before plotting.

### Cox regression and PDAC tumor grade regression

TCGA transcriptome counts and their associated clinical data were accessed using TCGAbiolinks (2.18.0). Downstream visualizations of the TCGA data were carried out on the VST transformed counts. The single grade 4 sample was combined with the grade 3 classification. To determine marginal survival associations between expression and survival, the survival status of the patients was regressed on the VST counts for each gene. The approximate z-score of each gene was computed using the estimated hazard and its standard error. To examine the association between gene expression and the observed grade of the tumor, we first omitted grade 1 (n=5) and grade 4 (n=1) tumors due to low sample sizes and focused on the grade 2 and 3 tumors. Then, we used a logistic regression model to evaluate if the expression of a gene was associated with a higher tumor grade. Estimated odds ratios were then standardized using their standard errors. To assess and visualize the distribution of these coefficients within the context of our RNA-seq clusters, the empirical cumulative distribution function (ECDF) of these scores were plotted with a 95% confidence envelope. To assess the medians of these scores within each RNA-seq cluster, a 95% percentile bootstrap was carried out on the median statistic with 10000 replications.

### PDAC tumor microarray analysis and GSEA

Pre-processed microarray data were accessed using the GEOquery (2.58.0) package. Pre-normalized values were used for each project and were subsequently analyzed using LIMMA (Linear Models for Microarray Data, 3.46.0). A paired subjects design was used to compare the matched adjacent normal tissue for each patient to the recorded tumor sample. Probe IDs were matched to genes according to the platform. In the case that a probe set contained multiple probes mapping to the same gene, the probe with the maximal average signal intensity within that probe set was used as the representative probe for that gene. To estimate the average fold changes across different studies, fold changes and standard errors reported by LIMMA were analyzed with a random effects meta-analysis design with Metafor (3.4.0). The pooled fold changes and standard errors were used to rank each gene according to the computed z-scores. The top 300 upregulated and 300 downregulated genes in *KCP* vs *GFP* PO samples were selected based on the fold change reported, among which 166 upregulated genes and 146 downregulated genes in *KCP* PO possessed a matching validated probe in the aforementioned meta-analysis. A GSEA was performed using fGSEA (1.16.0) based on the ranking of the genes in the meta-analysis to identify significant enrichments of our PO genes in the clinical samples. Assessment of significant enrichment was determined by normalized enrichment score and false discovery rate.

### ChIP-seq and analysis

PP1 and PP2 cells were resuspended and cross-linked with in PBS with 1% formaldehyde (Sigma, F1635) at room temperature for 15 minutes under rotation and quenched with 0.125 M glycine for 5 minutes at room temperature. Fixed cells were spun down and washed twice in cold PBS. Cell pellets were obtained by centrifugation at 3,000 RPM for 5 minutes at 4 °C and frozen in liquid nitrogen immediately before transferring to the −80 °C freezer. ∼10 million cells were used for one ChIP reaction. Cell pellets were thawed on ice, resuspended in 1 ml SDS lysis buffer (1% SDS, 10 mM EDTA, 50 mM Tris-HCl pH 8) containing proteinase and phosphatase inhibitor (Cell Signaling Technologies, #5872S) in an Eppendorf tube and incubated on ice for 10 minutes. Sonication was performed on a Branson 250 Sonifier with a 30% amplitude setting for 5.5 minutes (10 seconds on/off pulsing). Sonication products were spun down at 14,000 RPM at 4 °C for 10 minutes and 1 ml supernatant containing chromatin and DNA were transferred to a falcon tube containing 9 ml ChIP dilution buffer (0.01% SDS, 1.1% TritonX-100, 1.2 mM EDTA, 16.7 mM Tris-HCl pH 8, 167 mM NaCl) with proteinase inhibitor (cOmplete Protease inhibitor, 05056489001) and phosphatase inhibitor (Thermal Fisher, Scientific, 78427). 50 μl of Dynabeads (Life Technologies, 10009D) were added to samples and incubated at 4 °C with rotation for 1 hour. After pre-clearing, Dynabeads beads were removed and 200 μl of sample were collected as 2% input separately. 2.5 μg of antibodies were added to the pre-cleared samples for overnight incubation at 4 °C with rotation. 200 μl Dynabeads were added into one ChIP reaction and incubated for 4–6 hours at 4 °C with rotation. Dynabeads were collected by centrifugation with 3,000 RPM at 4 °C for 5 minutes and washed in 1 ml low salt buffer (0.1% SDS, 1% TritonX-100, 2 mM EDTA, 20 mM Tris-HCl pH 8, 150 mM NaCl) for 5 minutes at 4 °C with rotations. Then beads were washed in 1 ml high salt buffer (0.1% SDS, 1% TritonX-100, 2 mM EDTA, 20 mM Tris-HCl pH 8, 500 mM NaCl) twice and TE buffer (10 mM Tris-HCl pH 8, 1 mM EDTA) twice for 5 minutes at 4 °C with rotation. After the last wash, beads were resuspended in 250 μl elution buffer (1% SDS, 0.1 M NaHCO3) and incubated in a thermomixer: 850 RPM for 15 minutes at 60 °C. Supernatant were collected and 8ml of 5M NaCl was added for overnight decrosslinking at 65 °C. 10 μl 0.5 M EDTA, 20 μl 1 M Tris-HCl pH6.5 and 1 μl proteinase K (20 mg/ml) were added to de-crosslinked product and incubated for 1 hour at 45 °C. DNA was isolated using ChIP DNA Clean & Concentrator kit (Zymo, D5205). Libraries were prepared using NEBNext Ultra II DNA Library Prep Kit (NEB, E7645S) following manufacturer’s size selection and library preparation protocol.

ChIP-seq reads were first trimmed with fastp (0.23.4) and then aligned with Bowtie2 (2.4.1). Peak calling was performed by MACS2, with filterdup -auto and a p-value cutoff of 0.01. To generate associated tornado plots, Deeptools (3.5.1) was used with pileup tracks from MACS2. In the case IDR peaks were reported, MACS2 was run on the merged replicates for an experiment prior to generate a consensus peak set for that replicate. IDR was then performed with two replicates against the consensus peak set from the merged library.

### scRNA-seq analysis

Patient PDAC fastq files were downloaded from ^45^ and aligned with Cell Ranger count (Version 7.0.1). Per best practices of Seurat package (Version 4.1.0), the quality control of cells included 250 - 4000 as the filter for the number of features found in a cell and 0% - 20% as the filter for the percentage of tags mapped to the mitochondrial chromosome. Top 2000 variable genes and top 50 PCs were used for UMAP generation. Resolution = 0.15 was used for the initial call of FindClusters function. FindMarkers function was called to identify top expressed genes in each cluster. Cells were annotated based on the expression of marker genes: ductal cell 1 (*AMBP, CFTR, CA2),* ductal cell 2 (*KRT19, KRT7, MUC1, SLPI*), acinar cells *(PTF1A, CPA1, PRSS1, CTRB1, CTRB2, REG1B*), endocrine cell (*CHGB, CHGA, INS, IAPP),* stellate cell (*RGS5, ADIRF, PDGFRB, ACTA2),* fibroblast *(LUM, PDGFRA, DCN, COL1A1),* endothelial cell *(CDH5, CD34, PLVAP, VWF, CLDN5*), macrophage (*CD14, CD68, AIF1, FCGR1A*), T cell (*CD3E, CD3D, CD8A*), B cell (*CD19, CD79A, CD79B, MS4A1*). Ductal cell 1 and 2 were isolated for further analysis. Ductal cell only UMAP were generated by using top 2000 variable genes and top 50 PCs from ductal cells. Ductal cells were re-clustered and annotated as *MUC1* low, MUC1 intermediate and MUC1 high groups, representing normal, abnormal, and malignant epithelium population in patient PDAC samples as described ^45^, and validated by the increase of *FXYD3* and decrease of *AMBP* and *FXYD2*. Gene expression data extracted from the “data” slot, which was normalized and LN transformed, were used in the assay of the Seurat object for downstream analysis and violin plot. Model-based Analysis of Single-cell Transcriptomics package (MAST, 1.26.0) was applied to determine the fold change of gene expression among the identified clusters and statistical significance. The fold change of the average values across clusters was Log2 transformed. The p values were calculated by Wald tests after maximum likelihood estimation.

### Methyl capture and analysis

DNA methyl capture sequencing was performed on *KC, KP, KCP* and *KCPS* POs as previously described ^43^. POs in culture were directly lysed and gDNA was extracted using the DNeasy Blood & Tissue Kit (Qiagen, 69504) following the manufacturer’s guidelines. Two biological replicates for each genotype were submitted to the MSKCC Integrated Genomics Operation Core to be processed with Illumina’s TruSeq Methyl Capture EPIC Library Prep Kit and then sequenced. After PicoGreen quantification and quality control by Agilent BioAnalyzer, 480 - 490ng of gDNA were sheared using a LE220-plus Focused-ultrasonicator (Covaris catalog # 500569). Sequencing libraries were prepared using the KAPA Hyper Prep Kit (Kapa Biosystems KK8504) without PCR amplification. Post-ligation cleanup proceeded according to Illumina’s instructions with 110 μL Sample Purification Mix from the TruSeq Methyl Capture EPIC LT Library Prep Kit (Illumina catalog # FC-151-1002). After purification, 4 samples were pooled equimolar, and methylome regions were captured using EPIC oligos. Capture pools were bisulfite converted and amplified with 11 cycles of PCR. Pools were sequenced on a NovaSeq 6000 in a PE100 run using the NovaSeq 6000 S4 Reagent Kit (Illumina). The average number of read pairs per sample was 72 million.

Methyl capture data was aligned to the bisulfite-converted hg19 reference genome using Bismark v0.15.0 ^76^. Methylation status was extracted with the bismark_methylation_extractor script in Bismark. Replicates (n = 2) of each condition were merged. Differentially methylated regions (DMRs) between conditions were defined as regions containing at least five differentially methylated CpGs (DMCs; Chi-squared test followed by a Benjamini-Hochberg FDR procedure, BH-FDR < 20%, fold change > 1.25) and whose total methylation difference was more than 10% ^77,78^. To assess the methylation patterns over the ATAC-seq peak clusters, smoothed metaplots were manually constructed using mgcv (1.8-40). For each peak in the ATAC-seq cluster, the bigwig coverage track was used to assign the amount of DNA methylation within a 3000 bp window of the summit of the ATAC-seq peak. Afterward, the region around the peak was discretized into 100 bp bins. An additive model was fit to the signal as a function of the bin to generate smoothed signal tracks. The fitted data was then plotted for each group to compare the average methylation level within each ATAC-seq cluster.

### ATAC-seq of organoids

POs were isolated from Matrigel using Cell Recovery Solution, dissociated by accutase into single cells and re-suspended in PBS for Omni-ATAC-seq as previously described ^33,79^. ∼50,000 single cells were lysed in 100 ml ATAC-RSB lysis buffer (10 mM NaCl, 10 mM Tris pH7.4, 3 mM MgCl_2,_ 0.1% NP-40, 0.1% Tween-20, 0.01% Digitionin) on ice for 5min, washed in 1 ml cold wash buffer (10mM NaCl, 10mM Tris pH7.4, 3 mM MgCl_2_, 0.1% Tween-20). Nuclei were pelleted and re-suspended in 50 ml of transposition mix containing 1x TD Buffer (Illumina Cat #FC-121-1030), 2.5 µl Transposase (Illumina Cat #FC-121-1030), 16.5 µl PBS, 0.01% Digitonin, 0.1% Tween-20 and incubated in 37°C for 30 min. Reaction was cleaned up using a DNA Clean and Concentrator-5 Kit (Zymo, D4014). Libraries were prepared using the NEBNext Q5 Hot Start HiFi PCR Master Mix (NEB, M0543L) and Nextera primers. Prepared libraries were sent to MSKCC Integrated Genomics Operation core (IGO) and sequenced on a HiSeq 4000 sequencer in a PE50 run, using the HiSeq 3000/4000 SBS Kit (Illumina) or a NextSeq 500 sequencer in a PE50 run, using the TG NextSeq 500/550 High Output Kit v2.5 (150 Cycles) (Illumina). The runs yielded on average 35M reads per sample.

### ATAC-seq of archival human specimens

All patients were previously consented to MSKCC IRB#15-149 for research purposes. Archival patient PDAC samples from two 50 mm sections of OCT embedded tumors were broken up using razor blades in petri dish on ice in 1 ml NST-DAPI solution (116 mM NaCl, 8 mM Tris pH7.8, 0.8 mM CaCl_2_, 38 mM MgCl_2_, 0.4 mg/ml BSA, 5 mM EDTA, 0.16% NP-40 and 50 mg/ml DAPI) and filtered through top collection tube to remove debris. The crude nuclei preparation was then subjected to FACS on a BD FACSAria Flow Cytometer Cell Sorter to isolate 2N, 4N and aneuploid nuclei. Nuclei were collected in 200 µl of ATAC-RSB buffer (10 mM NaCl, 10 mM Tris pH7.8, 3 mM MgCl_2_) with 0.1% Tween-20 in 1.5ml Eppendorf tubes. Duplicates of sorted samples were used to confirm KRAS variant allele frequency (VAF). Approximately 10,000-50,000 nuclei from either aneuploid or 4N populations were carefully pelleted, resuspended in 50 ml of transposition mix and incubated in 37°C for 30 min. Reaction was cleaned up using Qiagen MinElute PCR Purification Kit followed by library preparation. Sequencing was performed on a HiSeq 4000 sequencer.

### ATAC-seq analysis

ATAC-seq reads were first checked with FastQC (0.11.9), trimmed with NGMerge (0.3) and then aligned with Bowtie2. MACS2 was used to identify peaks, and IDR was used to identify reproducible peaks at a cutoff of 0.01, which yields 73333 peaks for POs. Reads were then analyzed using DESeq2 (1.30.1). Differential peaks were defined by performing pairwise comparisons with a Wald Test, and then were reported at a BH adjusted p-value of 0.1. VST transformed counts were used for subsequent visualizations. Tornado plots and meta peaks were visualized using DeepTools (3.5.1). Clusters of peaks were identified as follows. First, DESeq2 was used to define all differentially accessible regions between the *GFP*, *KCP* and *KCPS* PO at an FDR cutoff of 0.1. Then the VST counts associated with those differential events were hierarchically clustered with correlation as the distance metric to identify 4 distinct clusters. To visualize PO samples and patient ATAC-seq data, the VST transformed counts were embedded in the same PCA plot.

Motifs were assigned to each peak using FIMO and Cis-BP (1.02) at a p-value of 1e-2. To determine the relationship between peak accessibility and the presence of the motif, we first identified all the peaks with a motif. Quantile regression (𝜏 = 0.5) was used to assess differences in the median fold changes for peaks associated with and without the motif. A positive coefficient from this regression indicates that the median fold change in the peaks with the motif are higher than the peaks without the motif. Similarly, a negative coefficient indicates that the median fold change in the peaks without the motif are higher than the ones with the motif. The associated t-statistic from this regression was used along with the odds ratio for differential accessibility in the associated direction and the proportion of peaks containing that motif to assess the strength of the relationship between the presence of a motif and peak opening or closing.

### CUT&RUN assay and analysis

CUT&RUN (C&R) was performed as previously described ^80^. 10 μl of Concanavalin A (Con A)-coated magnetic beads (Bangs Laboratories, BP531) were washed three times with 100 μl cold binding buffer (20 mM HEPES pH 7.9, 10 mM KCl, 1 mM CaCl2, 1 mM MnCl2). Activated Con A beads were resuspended in 10 μl cold binding buffer. 0.5M cells were harvested per C&R assay and were washed three times with 100 μl wash buffer (20 mM HEPES pH 7.5, 150 mM NaCl, 0.5 mM Spermidine, 1x protease inhibitor) at room temperature. Cells were resuspended in 100 μl wash buffer, mixed with 10 μl activated Con A beads and incubated for 10 min at room temperature. Cell permeabilization and primary antibody binding was performed by applying primary antibody at 1:100 dilution in 100 μl of cold antibody buffer (20 mM HEPES pH 7.5, 150 mM NaCl, 0.5 mM Spermidine, 0.01% Digitonin, 2 mM EDTA, 1x protease inhibitor cocktail) and incubating at room temperature for 20 min on a nutator. After primary antibody incubation, beads were washed three times with 1ml cold digitonin buffer (20 mM HEPES pH 7.5, 150 mM NaCl, 0.5 mM Spermidine, 0.01% Digitonin, 1x protease inhibitor cocktail). Beads were then resuspended with 100 μl cold digitonin buffer supplied with pAG-MNase (EpiCypher, 15-1016) followed by incubation at room temperature for 20 min on a nutator. Unbound pAG-MNase was removed by washing in 1 ml cold digitonin buffer two times. Beads were resuspended with 100 μl cold digitonin buffer and pAG-MNase was activated by reconstituting 2 mM CaCl2. Targeted chromatin cleavage was carried out by incubation at 0∼4°C for 30 min on a nutator. The reaction was stopped by adding 100 μl cold 2x stop buffer (340 mM NaCl, 20 mM EDTA, 4 mM EGTA, 0.01% Digitonin, 50 μg/ml RNase A). Cleaved chromatin was released by incubating at 37°C for 20 min on ThermoMixer at 800 rpm. C&R DNA was extracted from the supernatant using Monarch® PCR & DNA Cleanup Kit (NEB, T1030S). C&R library was prepared with no more than 10 ng C&R DNA using NEBNext Ultra II DNA Library Prep Kit (NEB) according to manufacturer’s protocol. For peak calling, C&R reads were processed using CUT&RUN Tools (1.0). For transcription factor C&R experiments, deduplicated reads whose insert spanned less than 120 bp were analyzed. For histone C&R, deduplicated reads whose insert spanned greater than 120 bp were analyzed. To generate associated tornado plots, Deeptools (3.5.1) was used as before with MACS2 pileup files generated using a library normalization.

## Supplemental information

Document1 Supplemental Figures 1-7 with figure legends

Table S1 Histology summary of xenograft assay

Table S4 Summary of patient PDAC samples used for ATAC-seq (related to Figure 3)

Table S5 Summary of top variable ATAC-seq peaks in all genotypes of POs (related to Figure 3)

---

**Figure S1.**
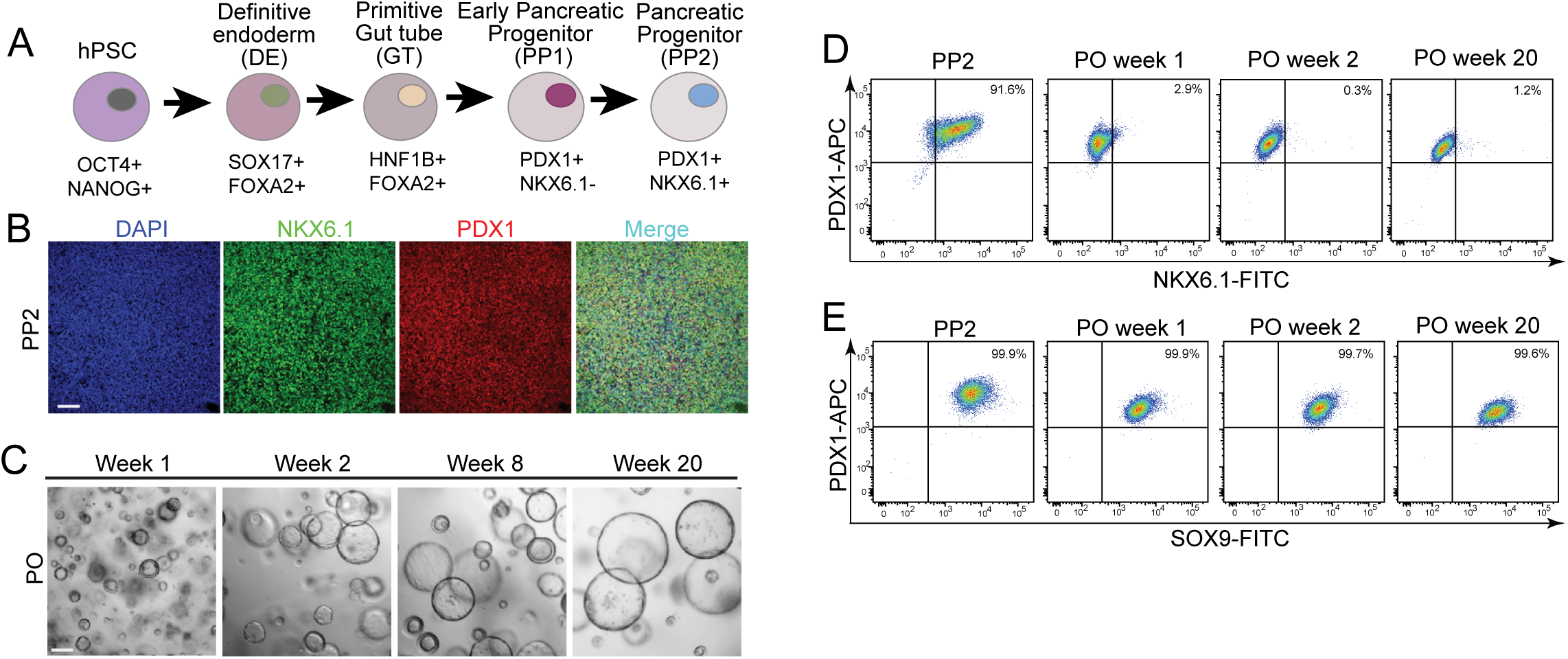
Establishment of an organoid culture enabling long-term propagation of pancreatic progenitors. (A) Scheme depicting the procedure of directed differentiation of hPSC into definitive endoderm (DE), primitive gut tube (GT), early pancreatic progenitor (PP1) and pancreatic progenitor (PP2) based on an optimized protocol. (B) Representative immunofluorescence image of PDX1/NKX6.1 in PP2 cell in 2D culture. Confocal images were maximum projected. Scale bar: 100μm. (C) Demonstration of long-term expansion of pancreatic progenitor organoid (POs) in culture. Scale bar: 100μm. (D,E) Flow cytometry analysis confirming the sustained expression of crucial pancreatic transcription factors PDX1/SOX9 and the loss of NKX6.1 in PO culture.

**Figure S2.**
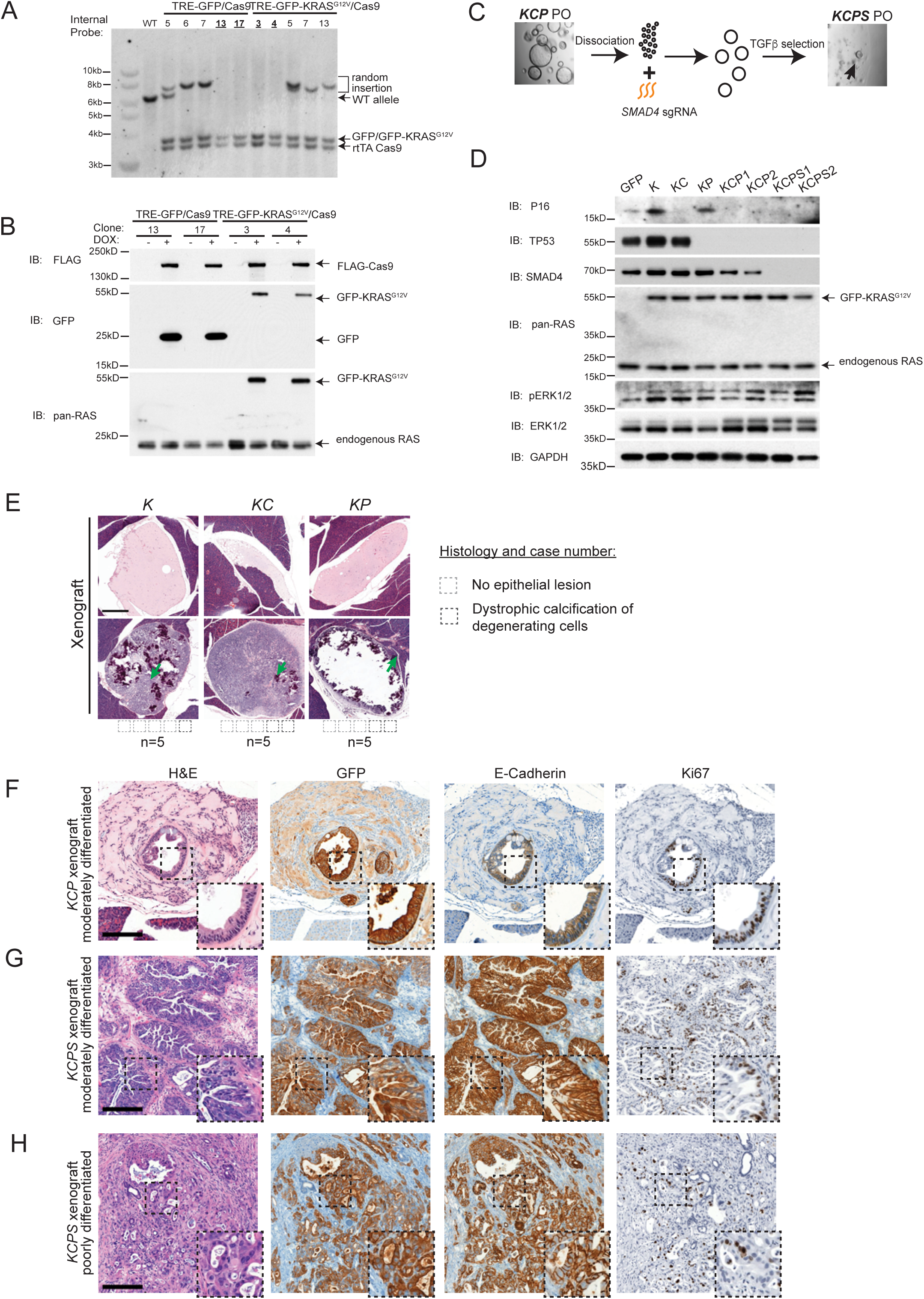
Validation of gene editing in POs and histology of xenografts. (A) Southern blot analysis of hPSC clones following insertion of GFP-*KRAS^G12V^*/*Cas9* or *GFP/Cas9* expression cassettes. An internal probe specifically recognized a 6492bp DNA fragment in wild-type genome, while presence of 3781bp and 3519bp fragments indicated successful insertion of *GFP* or *GFP*-*KRAS^G12V^* and *Cas9* cassettes. Clones exhibiting additional fragments indicated random insertion. (B) Southern blot verified hPSC clones (clone 13, 17 of *GFP/Cas9* lines and clone 3,4 of *KRAS^G12V^/Cas9* lines) were treated with doxycycline and examined by Western blot to confirm the expression of *GFP/GFP-KRAS^G12V^* and *Cas9* at the protein level. (C) The diagram illustrated the generation of *KCPS* POs through direct delivery of *SMAD4* sgRNA in *KCP* PO followed by TGF-b selection. CRISPR sequencing was performed in isolated PO clones. (D) Western blot confirmed the genotype of all POs established. (E) Mouse pancreata transplanted with *K*, *KC*, and *KP* POs were subjected to H&E staining. Occasional degenerating tissue with dystrophic calcification indicated by purple crystal in H&E staining (arrow) was observed at the injection site, scale bars: 500μm. (F-H) Moderately differentiated PDAC derived from *KCP* (F) and *KCPS* PO (G) and poorly differentiated PDAC from *KCPS* PO (H) were subject to H&E and IHC to visualize the expression of GFP-KRAS^G12V^, E-cadherin and Ki67 in tumor epithelia. Representative of 3 tumors, scale bars: 200μm.

**Figure S3.**
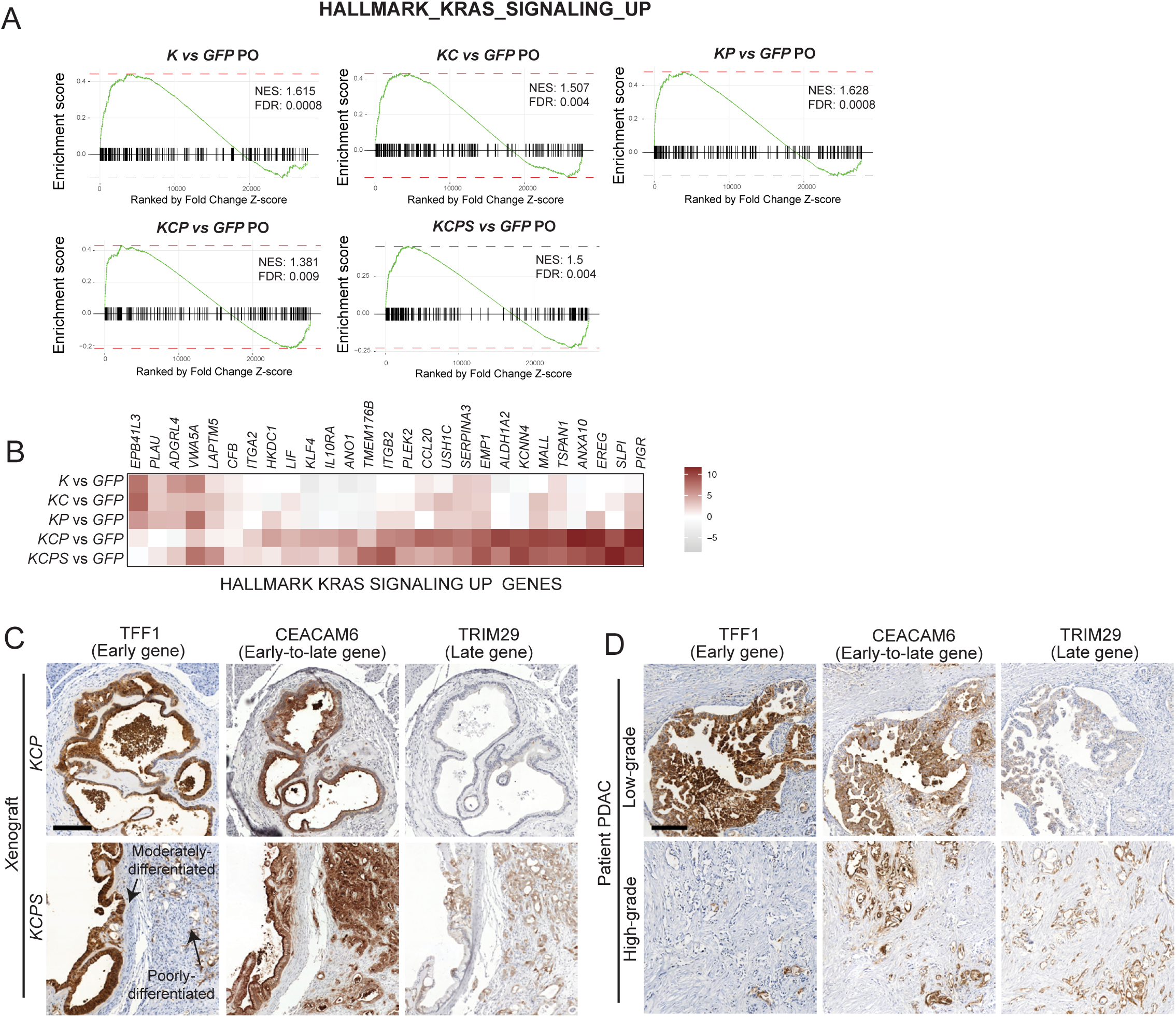
GSEA confirms the transcriptome signature of *KRAS* activation in POs and IHC validates the identified markers of early and late progression of PDAC. (A) Transcriptome profiles of *K*, *KC*, *KP*, *KCP*, and *KCPS* were compared with *GFP* POs in Gene Set Enrichment Analysis (GSEA). Significant enrichment of feature of HALLMARK KRAS SIGNALING UP gene set in *KRAS^G12V^* expressing POs comparing to that in *GPF* POs was observed. Normalized enrichment score (NES) and FDR values were presented. (B) The expression level of selected genes in HALLMARK KRAS SIGNALING UP set in *KRAS^G12V^* expressing POs was compared to that in *GPF* POs. Log2 fold changes were presented. (C-D) IHC assessment of the expression of additional early, early-to-late and late genes *TFF1*, *CEACAM6* and *TRIM29* in *KCP* and *KCPS* xenograft (C) or in low-grade and high-grade (D) human PDAC sections. Representative of 5 tumors each, scale bars: 200μm.

**Figure S4.**
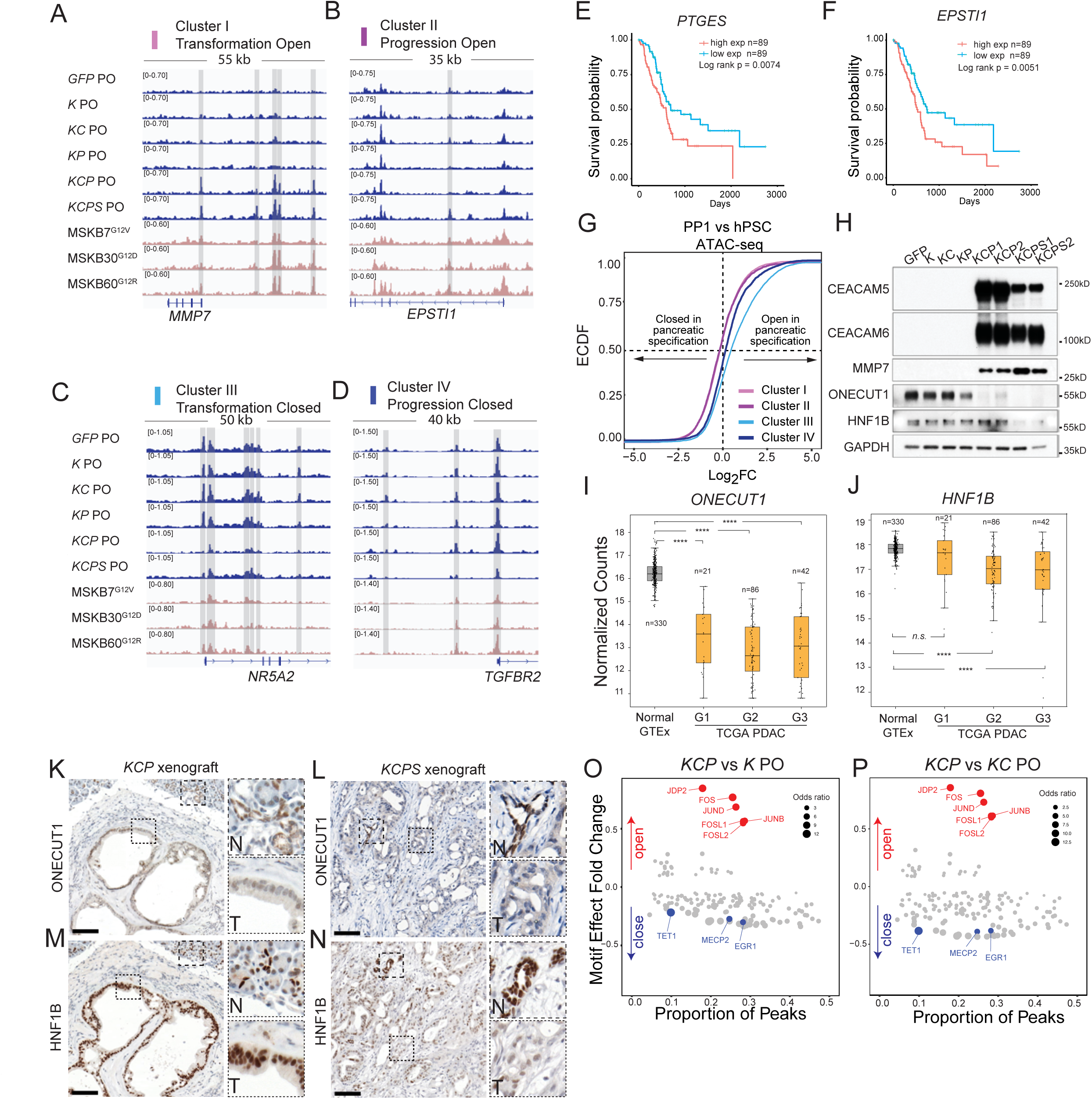
Chromatin accessibility profiling reveals two integral components of cellular plasticity during PDAC development. (A-D) Genomic visualization of representative loci associated with each of the clusters in POs defined in Figure 3C. Highlighted regions indicated identified peaks in each cluster in PO samples. (E,F) Kaplan Meier survival analysis was conducted based on TCGA PAAD dataset. (G) Chromatin accessibility changes during the differentiation of hPSC to PP1 was analyzed and outlined in an empirical cumulative distribution function (ECDF) plot. ATAC-seq signal fold change of PP1/hPSC was presented, and bootstrap test was applied to construct the 95% confidence intervals for the median of fold change for each cluster. The results were as follows: Cluster I: median Log_2_FC= -0.16, 95%CI [-0.18, -0.14]; Cluster II: median Log_2_FC= -0.22, 95%CI [-0.27, - 0.22]; Cluster III: median Log_2_FC= 0.62, 95%CI [0.56, 0.66]; Cluster IV: median Log_2_FC= 0.24, 95%CI [0.17, 0.30]. (H) Western blot validated the upregulation of Cluster I genes *CEACAM5/6* and *MMP7* and stepwise downregulation of Cluster III/IV gene *ONECUT1* and *HNF1B* during PDAC development. Same samples were examined as in Supplemental Figure 2D. (I,J) Analysis of *ONECUT1* and *HNF1B* expression in GTEx and TCGA PDAC samples. TCGA PDAC tumors were grouped by tumor grades, and VST normalized counts were plotted. Kruskal-Wallis test with Dunn’s multiple comparison test was performed. N.S. Padj>0.05, **** Padj<0.0001. (K-N) IHC analysis of *ONECUT1* and *HNF1B* expression in *KCP* and *KCPS* xenografts. Top insets (N) showed normal duct as a reference for staining. Lower insets (T) showed details of tumor epithelia. Representative of 5 tumors each, scale bars: 200μm. (O,P) Motif analysis comparing *KCP* PO with *K* and *KC* POs were performed as in Figure 3K,L.

**Figure S5.**
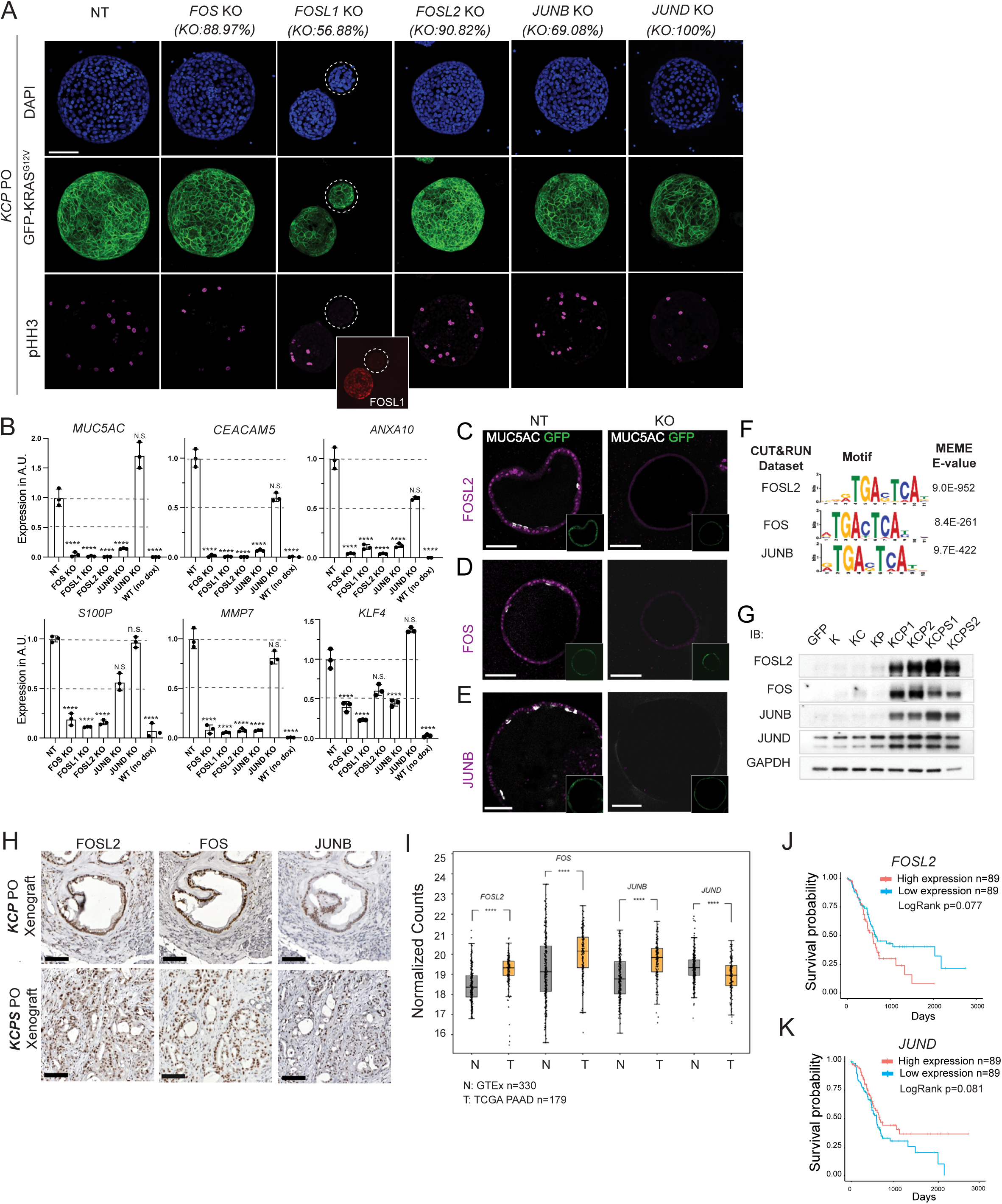
The acquisition of lineage plasticity during early PDAC transformation requires the activation of oncogenic AP-1 factors. (A) AP-1 perturbation was conducted in *KCP* POs and targeted CRISPR sequencing was performed four weeks after introducing sgRNAs. Immunofluorescence signal of pHH3 was visualized using confocal microscopy. Additional FOSL1 staining was performed to identify *FOSL1* knockout organoids in the mixed pool. Maximum projected images were presented, scale bar: 100 μm. (B) RT-qPCR was performed to examine the expression of malignant markers of PDAC in AP-1 KO *KCP* POs. Mean±SD, n=3. Samples with a |log2FC|>1 comparing to NT sgRNA treated control and displayed a significantly changed expression based on one-way ANOVA were labeled as significantly changed. **Padj<0.01, *** Padj<0.001, **** Padj<0.0001. (C-E) Representative immunofluorescence images of *MUC5AC* in *FOSL2*, *FOS* and *JUNB* KO *KCP* PO in culture were shown. The expression of *KRAS^G12V^* in the POs was indicated by GFP. Scale bars: 100μm. (F) MEME motif analysis identified AP-1 motifs in the FOSL2, FOS and JUNB C&R sequences. (G) Western blotting evaluated the expression of *FOSL2*, *FOS* and *JUNB* and *JUND* in POs. Same samples were examined as in Supplemental Figure 2D. (H) IHC of FOSL2, FOS and JUNB was performed on *KCP* and *KCPS* xenografts. Representative of 5 tumor samples each. Scale bars: 200μm. (I) The expression of *FOSL2*, *FOS*, *JUNB* and *JUND* in GTEx (V8) and TCGA PAAD datasets was examined. Wilcoxon signed rank test was performed, **** p<0.0001. (J,K) Kaplan Meier survival analysis was performed based on TCGA PAAD dataset. p values of log-rank test were annotated.

**Figure S6.**
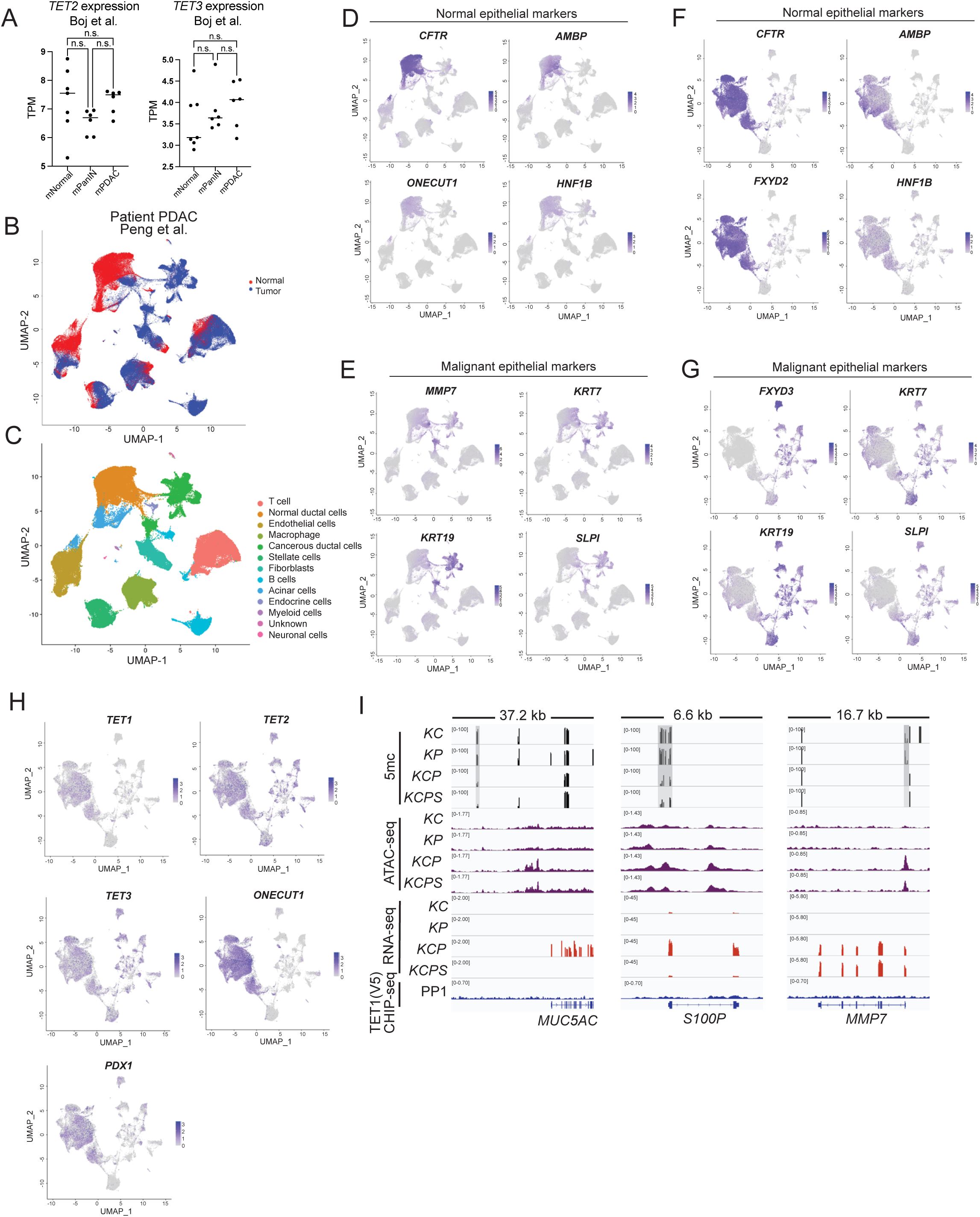
*TET1* loss coincides with the suppression of pancreatic lineage program. (A) Expression level of *TET2* and *TET3* in mouse normal duct, PanIN, and PDAC organoids from Boj et al. was displayed in TPM, one-way ANOVA. (B-H) A patient PDAC data set containing 24 patient PDAC and 11 normal tissue scRNA-seq samples was processed using Seurat single cell RNA-seq package. Initial clustering and marker finding function with known cell type markers allowed the identification of 10 major cell types mentioned in the original study: ductal cell 1 (normal duct), ductal cell 2 (cancerous duct), acinar cells, endocrine cell, stellate cell, fibroblast, endothelial cell, macrophage, T cell, B cell. Two additional cell types were annotated: myeloid cells and neuronal cells (B,C). Expression of normal epithelial markers *CFTR*, *AMBP*, *ONECUT1*, *HNF1B* and malignant epithelial markers *MMP7*, *KRT7*, *KRT19* and *SLPI* were verified by marker finding function and feature plots (D,E). Subsequent re-clustering of ductal cell 1 and ductal cell 2 populations, the major epithelial components in the data set, revealed 3 clusters based on *MUC1* expression levels: normal (*MUC1* low), abnormal (*MUC1* slightly increased), and malignant epithelia (*MUC1* high). Similar expression pattern was observed with malignant markers *FXYD3*, *KRT7*, *KRT19* and *SLPI* (G). Decreased expression of normal ductal makers *CFTR, AMBP, FXYD2* and *HNF1B* was observed in malignant epithelia (*MUC1* high) (F). *TET1*, *TET2*, *TET3*, *ONECUT1* and *PDX1* expression levels were examined on the feature plots (H). (I) Genomic visualization of methylation signals in *KC*, *KP*, *KCP* and *KCPS* POs was presented alongside ATAC-seq and RNA-seq tracks, for loci associated with Cluster I genes *MUC5AC*, *S100P* and *MMP7.* No TET1 bound regions were called in these loci.

**Figure S7.**
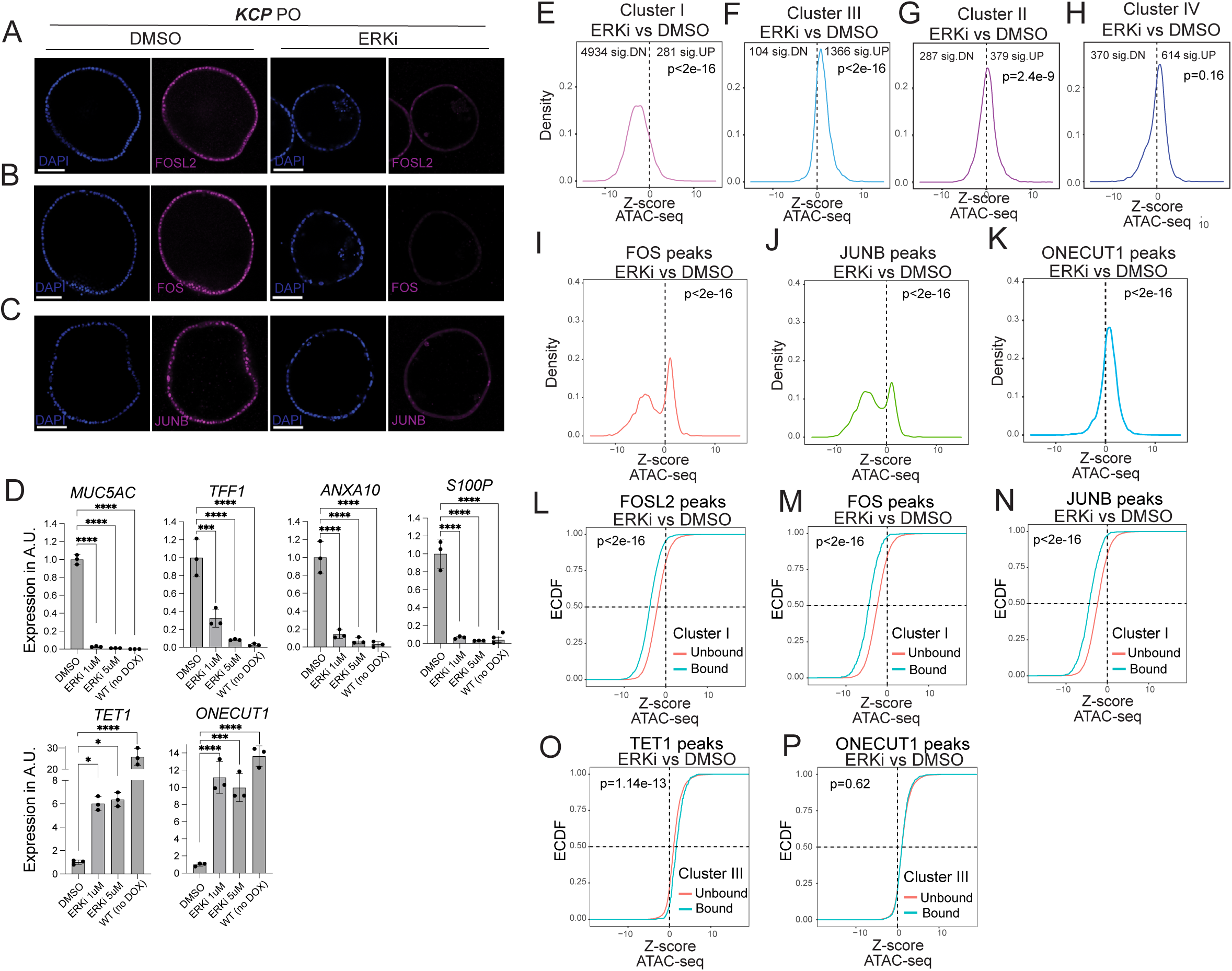
ERK activates oncogenic AP-1 and suppresses TET1-mediated pancreatic lineage program in early PDAC transformation. (A-C) Immunofluorescence assay demonstrated the impact of Temuterkib treatment on AP-1 protein levels in *KCP* POs. Scale bars: 100μm. (D) RNA was extracted from *KCP* POs treated with Temuterkib as in (A-C) for RT-qPCR analysis. A non-doxycycline treated PO served as wild-type control. Mean±SD, n=3, *Padj<0.05,**Padj<0.01, *** Padj<0.001, **** Padj<0.0001, one-way ANOVA. (E-H) Density plots showed the effect of Temuterkib treatment on ATAC-seq signal in Cluster I-IV regions in *KCP* POs (related to Figure 6C). Wilcoxon signed rank test was performed. (I-K) Density plots illustrated the changes in ATAC-seq signal of FOS, JUNB, and ONECUT1 bound regions in Temuterkib or DMSO-treated *KCP* POs (related to Figure 6E,F). Wilcoxon signed rank test was performed. (L-N) ECDF plots exhibited the ratio of ATAC-seq signal in FOSL2, FOS, and JUNB bound or unbound Cluster I regions in Temuterkib or DMSO-treated *KCP* POs. Wilcoxon signed rank test was performed. (O,P) ECDF plots showed the ratio of ATAC-seq signal of TET1 and ONECUT1 bound or unbound Cluster III regions in Temuterkib or DMSO-treated *KCP* POs. Wilcoxon signed rank test was performed.

**Table S1.**
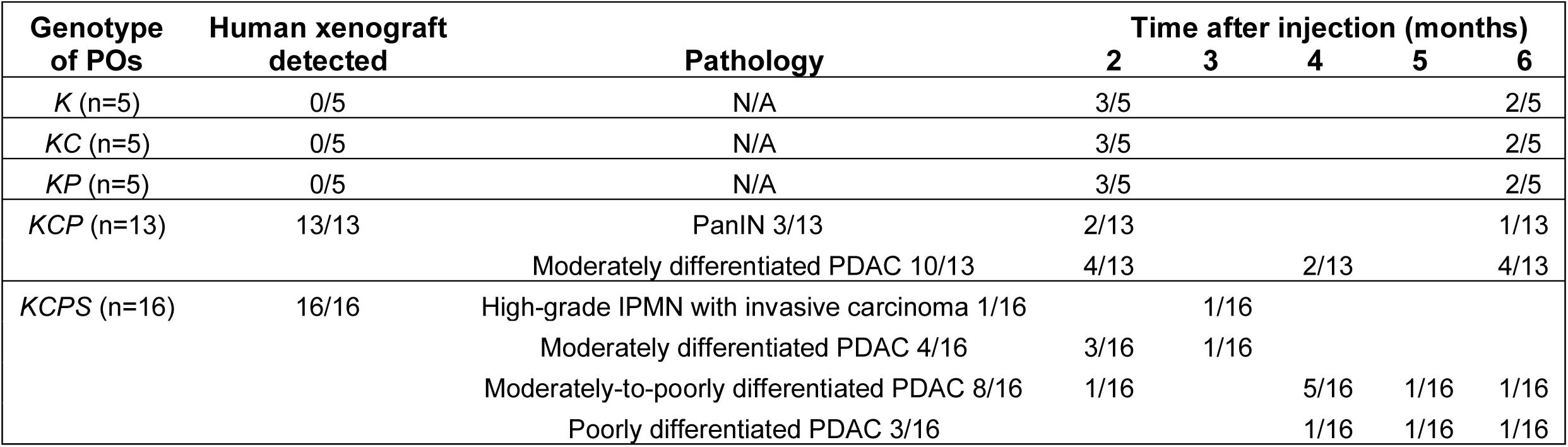
Summary of histology of all mouse xenograft samples analyzed.

| Genotype of POs | Human xenograft detected | Pathology | Time after injection (months) |  |  |  |  |
| --- | --- | --- | --- | --- | --- | --- | --- |
|  |  |  | 2 | 3 | 4 | 5 | 6 |
| *K* (n=5) | 0/5 | N/A | 3/5 |  |  |  | 2/5 |
| *KC* (n=5) | 0/5 | N/A | 3/5 |  |  |  | 2/5 |
| *KP* (n=5) | 0/5 | N/A | 3/5 |  |  |  | 2/5 |
| *KCP* (n=13) | 13/13 | PanIN 3/13 | 2/13 |  |  |  | 1/13 |
|  |  | Moderately differentiated PDAC 10/13 | 4/13 |  | 2/13 |  | 4/13 |
| *KCPS* (n=16) | 16/16 | High-grade IPMN with invasive carcinoma 1/16 |  | 1/16 |  |  |  |
|  |  | Moderately differentiated PDAC 4/16 | 3/16 | 1/16 |  |  |  |
|  |  | Moderately-to-poorly differentiated PDAC 8/16 | 1/16 |  | 5/16 | 1/16 | 1/16 |
|  |  | Poorly differentiated PDAC 3/16 |  |  | 1/16 | 1/16 | 1/16 |

**Table S2.** Summary of meta-analysis and top up and down-regulated genes in *KCP* vs *GFP* POs used for GSEA (related to Figure 2, spreadsheet attached)

**Table S3.** Summary of clustering of top 3000 variable genes across all genotypes of POs (related to Figure 2, spreadsheet attached)

**Table S4.**
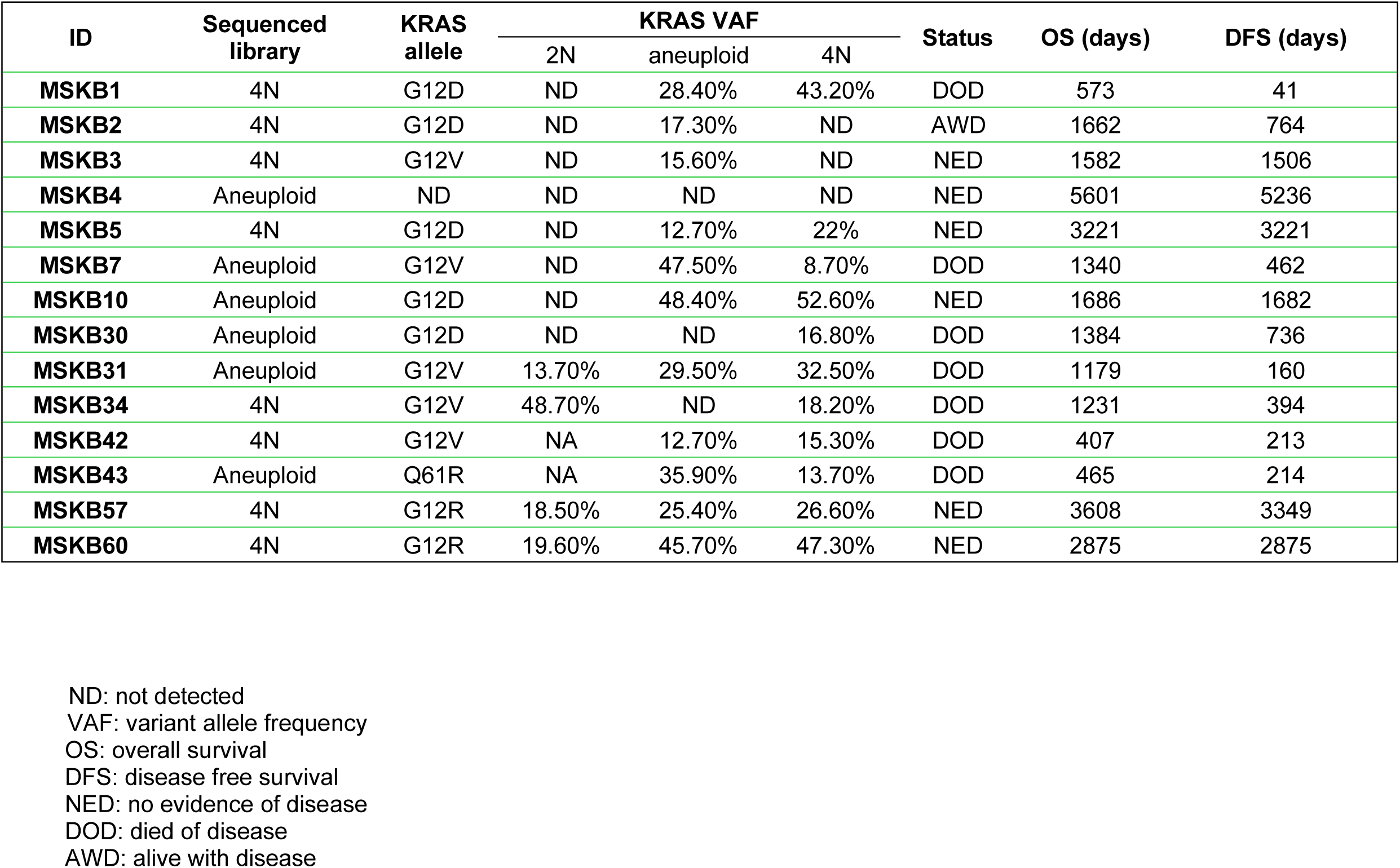
Summary of patient PDAC samples used for ATAC-seq.

| ID | Sequenced library | KRAS allele | KRAS VAF |  |  | Status | OS (days) | DFS (days) |
| --- | --- | --- | --- | --- | --- | --- | --- | --- |
|  |  |  | 2N | aneuploid | 4N |  |  |  |
| **MSKB1** | 4N | G12D | ND | 28.40% | 43.20% | DOD | 573 | 41 |
| **MSKB2** | 4N | G12D | ND | 17.30% | ND | AWD | 1662 | 764 |
| **MSKB3** | 4N | G12V | ND | 15.60% | ND | NED | 1582 | 1506 |
| **MSKB4** | Aneuploid | ND | ND | ND | ND | NED | 5601 | 5236 |
| **MSKB5** | 4N | G12D | ND | 12.70% | 22% | NED | 3221 | 3221 |
| **MSKB7** | Aneuploid | G12V | ND | 47.50% | 8.70% | DOD | 1340 | 462 |
| **MSKB10** | Aneuploid | G12D | ND | 48.40% | 52.60% | NED | 1686 | 1682 |
| **MSKB30** | Aneuploid | G12D | ND | ND | 16.80% | DOD | 1384 | 736 |
| **MSKB31** | Aneuploid | G12V | 13.70% | 29.50% | 32.50% | DOD | 1179 | 160 |
| **MSKB34** | 4N | G12V | 48.70% | ND | 18.20% | DOD | 1231 | 394 |
| **MSKB42** | 4N | G12V | NA | 12.70% | 15.30% | DOD | 407 | 213 |
| **MSKB43** | Aneuploid | Q61R | NA | 35.90% | 13.70% | DOD | 465 | 214 |
| **MSKB57** | 4N | G12R | 18.50% | 25.40% | 26.60% | NED | 3608 | 3349 |
| **MSKB60** | 4N | G12R | 19.60% | 45.70% | 47.30% | NED | 2875 | 2875 |

ND: not detected VAF: variant allele frequency OS: overall survival DFS: disease free survival NED: no evidence of disease DOD: died of disease AWD: alive with disease

**Table S5.** Summary of top variable ATAC-seq peaks in all genotypes of POs (related to Figure 3, spreadsheet attached)

**Table S6.** Summary of differentially expressed gene analysis in AP-1 knockout and ERKi treated *KCP* POs (related to Figure 4,6, spreadsheet attached)

**Table S7.**
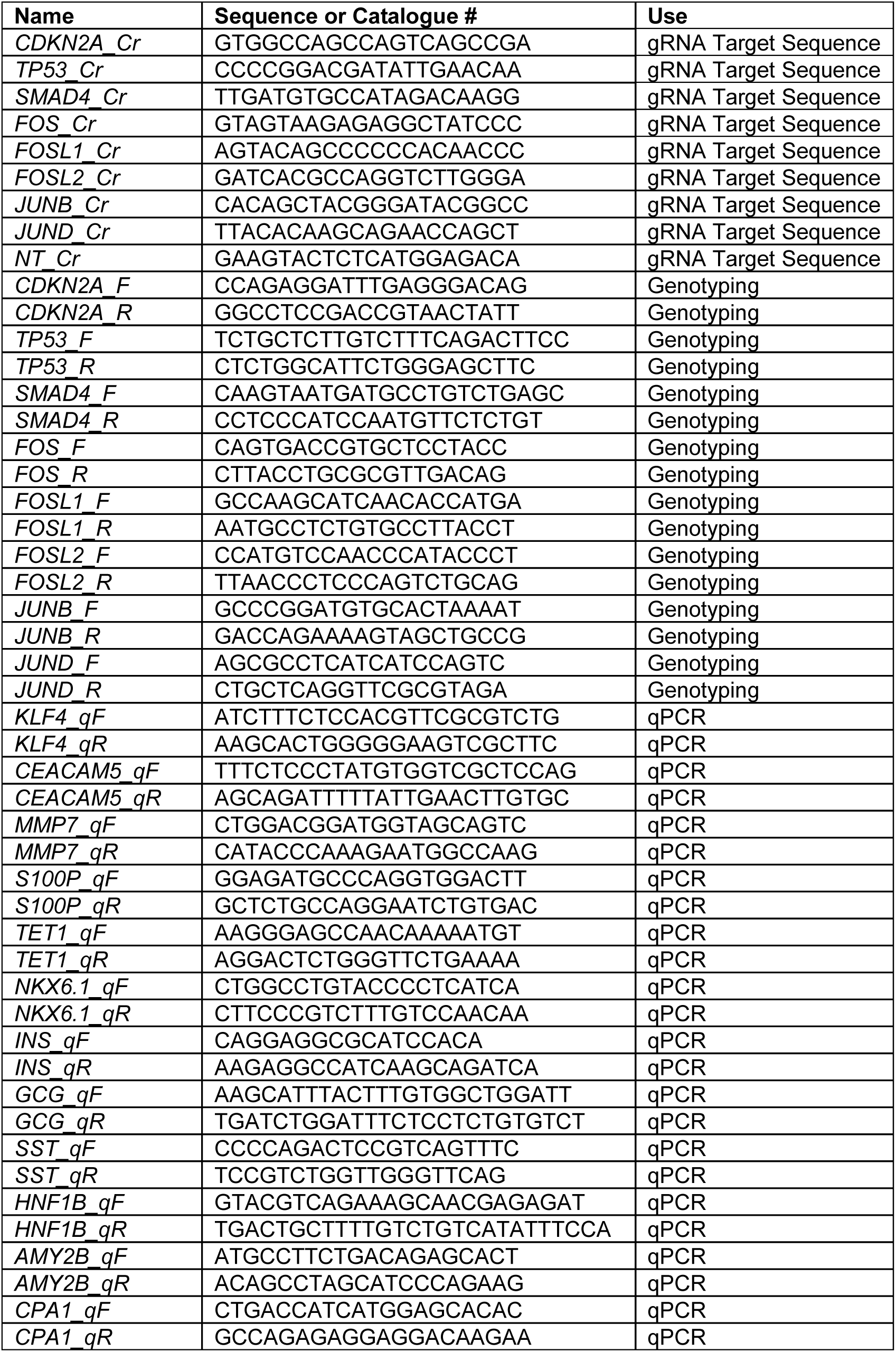
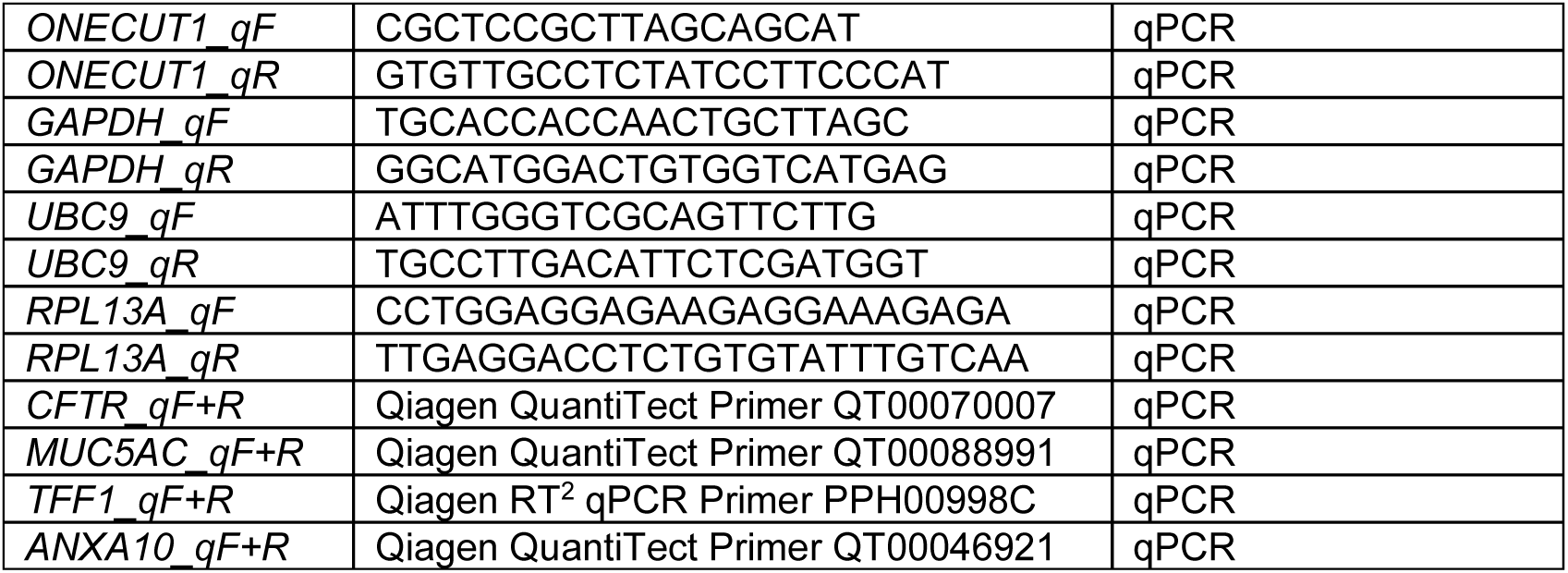
List of sgRNA target sequences, primer sequences for genotyping and RT-qPCR.

**Table S8.** List of primary antibodies. Antibody targets, company or lab that supplied antibodies, catalog numbers, applications, and quantity or dilution used are indicated.

| Name | Company/Lab | Cat# | Use (Dilution) |
| --- | --- | --- | --- |
| PDX1 | R&D Systems | AF2419 | Flow (1:500)<br>2D IF (1:500)<br>3D IF (1:200)<br>ChIP (5µg /10M cells) |
| NKX6-1 | DSHB | F55A12 | Flow (1:500)<br>2D IF (1:500)<br>3D IF (1:100) |
| SOX9 | Millipore | ab5535 | Flow (1:500) |
| FLAG (M2) | Sigma-Aldrich | F1849-200ug | WB (1:2000) |
| GFP (JL-8) | Takara Bio Clontech | 632381 | WB (1:500) |
| GFP | Abcam | ab13970 | IHC (2µg/ml) |
| Pan-RAS | Millipore | op40 | WB (1:500) |
| CDKN2A (p16 INK4A) | Cell Signaling Technologies | 80772 | WB (1:500) |
| TP53 (DO-1) | Santa Cruz | sc-126 | WB (1:1000) |
| SMAD4 | Cell Signaling Technologies | 38454 | WB (1:500) |
| ERK1/2 | Sigma-Aldrich | M5670-25UL | WB (1:10000) |
| Phosphor-ERK1/2 | Sigma-Aldrich | M8159-50UL | WB (1:3000) |
| GAPDH | Santa Cruz | sc-25778 | WB (1:2000) |
| E-Cadherin | BD Bioscience | 610181 | IHC (2.5µg/ml) |
| E-Cadherin | Invitrogen | 13-1900 | 3D IF (1:100) |
| Ki67 | ThermoFisher | 18-0191Z | IHC (1:200) |
| CEACAM5 | Sigma-Aldrich | HPA019758 | IHC (1:1000) |
| CEACAM5 | R&D Systems | MAB41281 | WB (1:2000) |
| MMP7 | R&D Systems | AF907 | IHC (1:750) |
| MMP7 | R&D Systems | MAB9071 | WB (1:1000) |
| PTGES | Sigma-Aldrich | HPA045064 | IHC (1:1200) |
| TFF1 | Sigma-Aldrich | HPA003425 | IHC (1:2000) |
| CEACAM6<br>(KOR-SA3544) | MBL | D028-3 | IHC (1:2000)<br>WB (1:2000) |
| TRIM29 | Sigma-Aldrich | HPA020053 | IHC (1:2000) |
| ONECUT1 | Santa Cruz | sc-13050 | ChIP (5µg/10M cells)<br>IHC (1:100)<br>WB (1:500) |
| HNF1B | Sigma-Aldrich | HPA002083 | IHC (1:2000) |
| HNF1B | BD Bioscience | 612504 | WB (1:500) |
| FOS | Cell Signaling Technologies | 2250 | WB (1:1000) |
| FOS | Sigma-Aldrich | HPA018531 | 3D IF (1:500)<br>IHC (1:1000)<br>CR (1:100) |
| FOSL1 | Santa Cruz | sc-28310 | 3D IF (1:200) |
| FOSL1 | Cell Signaling Technologies | 5281 | WB (1:300) |
| FOSL2 | Sigma-Aldrich | HPA004817 | WB (1:2000)<br>3D IF (1:500)<br>IHC (1:2000)<br>CR (1:100) |
| JUNB | Cell Signaling Technologies | 3753 | WB (1:500)<br>3D IF (1:200)<br>IHC (1:1000)<br>CR (1:100) |
| JUND | Cell Signaling Technologies | 5000 | WB (1:1000) |
| KRT19 | Millipore | mab3238 | IHC (1:1000) |
| MUC5AC | Abcam | ab3649 | 3D IF (1:300) |
| pHH3 | Cell Signaling Technologies | 9701 | 3D IF (1:200) |
| 5hMC | Active Motif | 39791 | IHC (1:30000) |
| V5-tag | Cell Signaling Technologies | 13202 | CHIP (5ug/10M cell) |

